# Recent evolutionary history of tigers highlights contrasting roles of genetic drift and selection

**DOI:** 10.1101/696146

**Authors:** Ellie E. Armstrong, Anubhab Khan, Ryan W Taylor, Alexandre Gouy, Gili Greenbaum, Alexandre Thiéry, Jonathan TL Kang, Sergio A. Redondo, Stefan Prost, Gregory Barsh, Christopher Kaelin, Sameer Phalke, Anup Chugani, Martin Gilbert, Dale Miquelle, Arun Zachariah, Udayan Borthakur, Anuradha Reddy, Edward Louis, Oliver A. Ryder, Y V Jhala, Dmitri Petrov, Laurent Excoffier, Elizabeth Hadly, Uma Ramakrishnan

## Abstract

Formulating strategies for species conservation requires knowledge of evolutionary and genetic history. Tigers are among the most charismatic of endangered species and garner significant conservation attention. However, the evolutionary history and genomic variation of tigers remain poorly known. With 70% of the worlds wild tigers living in India, such knowledge is critical for tiger conservation. We re-sequenced 65 individual tiger genomes across their extant geographic range, representing most extant subspecies with a specific focus on tigers from India. As suggested by earlier studies, we found strong genetic differentiation between the putative tiger subspecies. Despite high total genomic diversity in India, individual tigers host longer runs of homozygosity, potentially suggesting recent inbreeding, possibly because of small and fragmented protected areas. Surprisingly, demographic models suggest recent divergence (within the last 10,000 years) between populations, and strong population bottlenecks. Amur tiger genomes revealed the strongest signals of selection mainly related to metabolic adaptation to cold, while Sumatran tigers show evidence of evolving under weak selection for genes involved in body size regulation. Depending on conservation objectives, our results support the isolation of Amur and Sumatran tigers, while geneflow between Malayan and South Asian tigers may be considered. Further, the impacts of ongoing connectivity loss on the health and persistence of tigers in India should be closely monitored.

## Introduction

Species are classified as endangered based on recent trends in their population sizes and habitat quality (e.g. IUCN red list criteria). Endangerment status spurs funding, conservation action, and management in an attempt to secure species survival. Implicit assumptions underpinning risk category designations are that recent demographic trends determine extinction probability and that loss of genetic diversity and inbreeding in small populations compromise their fitness. Empirical, theoretical, and experimental studies also suggest that individual and population survival is contingent on genetic variability (Saccheri et al., 1998). Standing genetic variation in a population is determined by the interplay of mutation rate, demography, gene flow/connectivity, selection, and genetic drift over time (Ellegren & Galtier, 2016). For endangered species that are characterized by long-term decline, their small and fragmented populations, unique selection pressures, and more frequent mating between close relatives, may result in unique population histories resulting in low, but distinct standing genetic variation. If populations remain connected despite landscape fragmentation, and in the absence of differential selection, most standing genetic variation may be shared. Importantly, populations and landscapes within species distributions might have diverse histories, and hence differential probabilities of survival contingent on their standing genetic variation.

Genetic diversity is often used as a proxy for evolutionary divergence, usually without detailing whether differences result from adaptation to local environments, stochastic drift, or both. Such understanding has been elusive until recently because estimating recent history of populations requires large genomic sampling across populations for high statistical power and appropriate techniques for detection of recent selection across the genome (Pool et al., 2010). Recent advances in sequencing technology, growth of population genomic models, and enhanced computing power have revolutionized our ability to read and interpret genomes, thus allowing quantification of the sum total of genetic variation within individuals and populations.

For several endangered species, whole genome re-sequencing has revealed low species-level variation (e.g. Iberian lynx, Abascal et al., 2016), strong signatures of population decline (e.g. mountain gorillas: Xue et al., 2015) and recent inbreeding in isolated populations (wolves: Kardos et al., 2018). Genomic analyses have also identified mutations pre-disposing individuals to disease (Tasmanian devil: Murchison et al 2012) as well as recent protective mutations (Tasmanian devil: Epstein et al., 2016). Finally, genomics has identified signatures of population decline in extinct species (woolly mammoth: Palkopoulou et al., 2015) and strong signatures of selection prior to extinction (passenger pigeon: Murray et al. 2017).

Initial studies typically sequence high-coverage genomes of a few individuals, often from ex situ collections or voucher specimens, to infer levels of variation. To better understand population genetics of endangered species, genome sequencing efforts should be at larger geographic scales, sampling surviving populations comprehensively. Broader sampling is made particularly challenging in wide-ranging endangered species, especially those with geographic ranges spanning international borders, where both sampling permissions and population management strategies differ.

The tiger (*Panthera tigris*) is an iconic and charismatic endangered species that once spanned 70 degrees of latitude across Asia. Between 2,154 and 3,159 tigers remain, now occupying less than 6% of their 1900 A.D. range (Goodrich et al., 2015). Despite this recent range collapse, tigers still live across 11 Asian nations in an extraordinary range of habitats that include estuarine mangrove forests (the Sundarbans), dry deciduous forests (parts of India), tropical rainforests (Malay Peninsula) and cold, temperate forests (Russian Far East). However, the specific adaptations of the various populations to their habitats remain largely unknown.

Tigers have been classified into four extant (and four extinct) subspecies (Nowell and Jackson, 1996), although genetic and other data substantiated (e.g. Luo et al., 2004, suggested an additional population group) or contradicted (Wilting et al., 2015, suggested fewer population groups) this classification. Liu et al. (2018) presented the first analyses of genome-wide variation using voucher specimens across the tiger range, and inferred relatively old divergences (∼68,000 years ago) between subspecies with low subsequent gene flow (1-10%). However, their sampling of the most populous (Jhala et al. 2015) and genetically diverse tiger subspecies—the South Asian tiger— was limited across habitats.

To formulate effective conservation strategies for such widely distributed yet fragmented species, a comprehensive understanding of range-wide genomic variation, demographic history and adaptation, and recent impacts of fragmentation is needed. This must include genomes sampled from the various landscapes and populations across the range. Here, we use whole genomes from across the wild tiger range, with representation from four extant subspecies (except *Panthera tigris corbettii*) and most habitats to infer historical and recent evolutionary history of tigers by investigating (a) population clustering within range-wide samples, (b) genomic variation, (c) possible signatures of recent inbreeding, and (d) demographic history and differential selection. Such an approach, can provide insights on how, or whether, evolutionary trajectories of tiger populations should be maintained in the future.

## Results and Discussion

We sequenced genomes from 65 individuals (Figure 1A, Supplementary Table 1) at varying coverage (4.2X-32.9X, median 14.4X). Our samples included wild-caught and captive-bred tigers from four putative extant sub-specific regions (South Asia, Malayan peninsula, East Siberia and Sumatra). We were unable to sample the South China tiger *(P. t. amoyensis*), considered extinct-in-the-wild. While the South China tiger is thought to be ancestral, Liu et al. (2018) suggested uncertainty about the antiquity of this population, since mitochondrial genomes were similar to those of Amur tigers.

**Figure 1:**
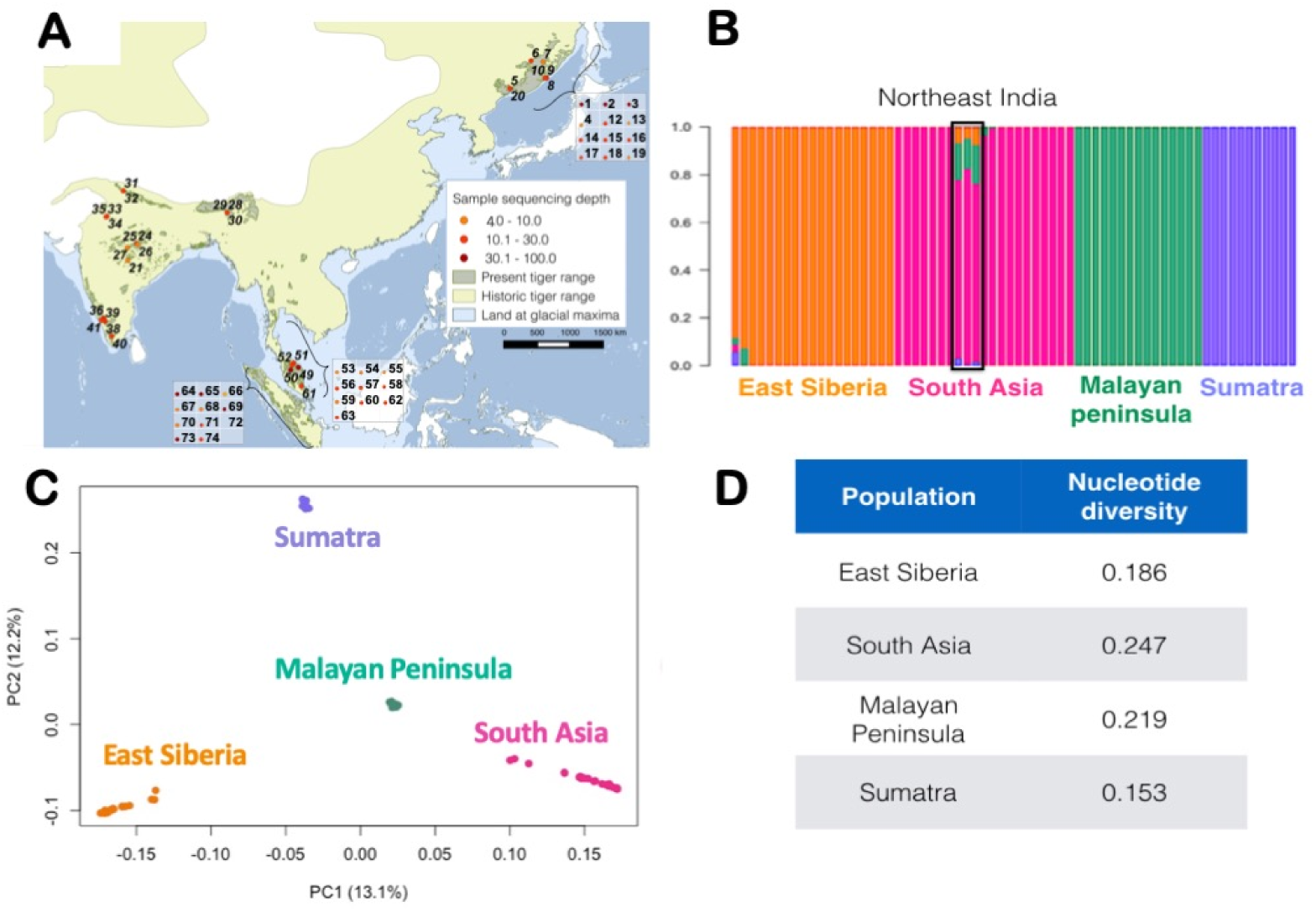
(A) Map of tiger samples used in this study. Each number refers to an individual. Genomic sequence coverage for each sample is colour coded. Wild samples (n=32) are represented on the map, while captive individuals (n=34) are in boxes. Sample details presented in Supplementary table 1. Historical and present range map courtesy IUCN (Goodrich et al 2015), (B) ADMIXTURE (K=4), (C) Principal component analyses (PCA) revealing genetic population structure in tigers. Colors in both ADMIXTURE and PCA analyses denote individuals from the different geographical regions and (D) nucleotide diversity (pi) estimates for tigers from different regions.

### Population structure

Model-based ADMIXTURE analyses suggested that genetically distinct populations are concordant with earlier definitions of subspecies (as also suggested by Luo et al., 2019 and Liu et al., 2018) (Figure 1B). Cross-validation statistics suggested that K=4 fit the data best (Supplementary Figure 2). At K=4, South Asian individuals sampled from the northeastern region of India show some admixture with Malayan individuals and to a lesser extent with Amur and Sumatran individuals (also see Figure 1B). At higher K (K=5, Supplementary Figure 3) the data reveal substructure within India separating south Indian tigers from others in India, but no further substructure in the other subspecies. Higher values of K fit the data poorly.

PCA (Figure 1C) revealed a similar pattern to the ADMIXTURE analyses, with the subspecies forming discrete clusters. PC1 and PC2 separate the four groups/subspecies (PC1: 13.1%; PC2:12.2%). PC1 separate the four groups in a north-to-south direction, while PC2 resolved individuals in the east-to-west direction. We henceforth refer to the geographic regions by their sub-specific names (East Siberia: Amur; South Asia: South Asian; Malay Peninsula: Malayan; and Sumatra: Sumatran). Additionally, PC1 shows stronger similarity between South Asian and Malayan tigers than South Asian and Sumatran tigers, consistent with the majority of results from K=3. PC2 (12.2% variation) and PC3 (10.9% variation) further separate the four groups, and also separated some individuals within populations (Supplementary Figures 4 and 6). In contrast, PCA analysis of non-transcribed regions including only high-coverage individuals (coverage > 20X) within the dataset (Sumatran=3; South Asian=3, Malayan=3, Amur=3, Supplementary Table 6) suggested that the Amur population is much less differentiated and closer to the Malayan population (Figure 3B). It is possible that both Amur and Malayan populations were close to a putative ancestral Asian tiger population.

**Figure 2:**
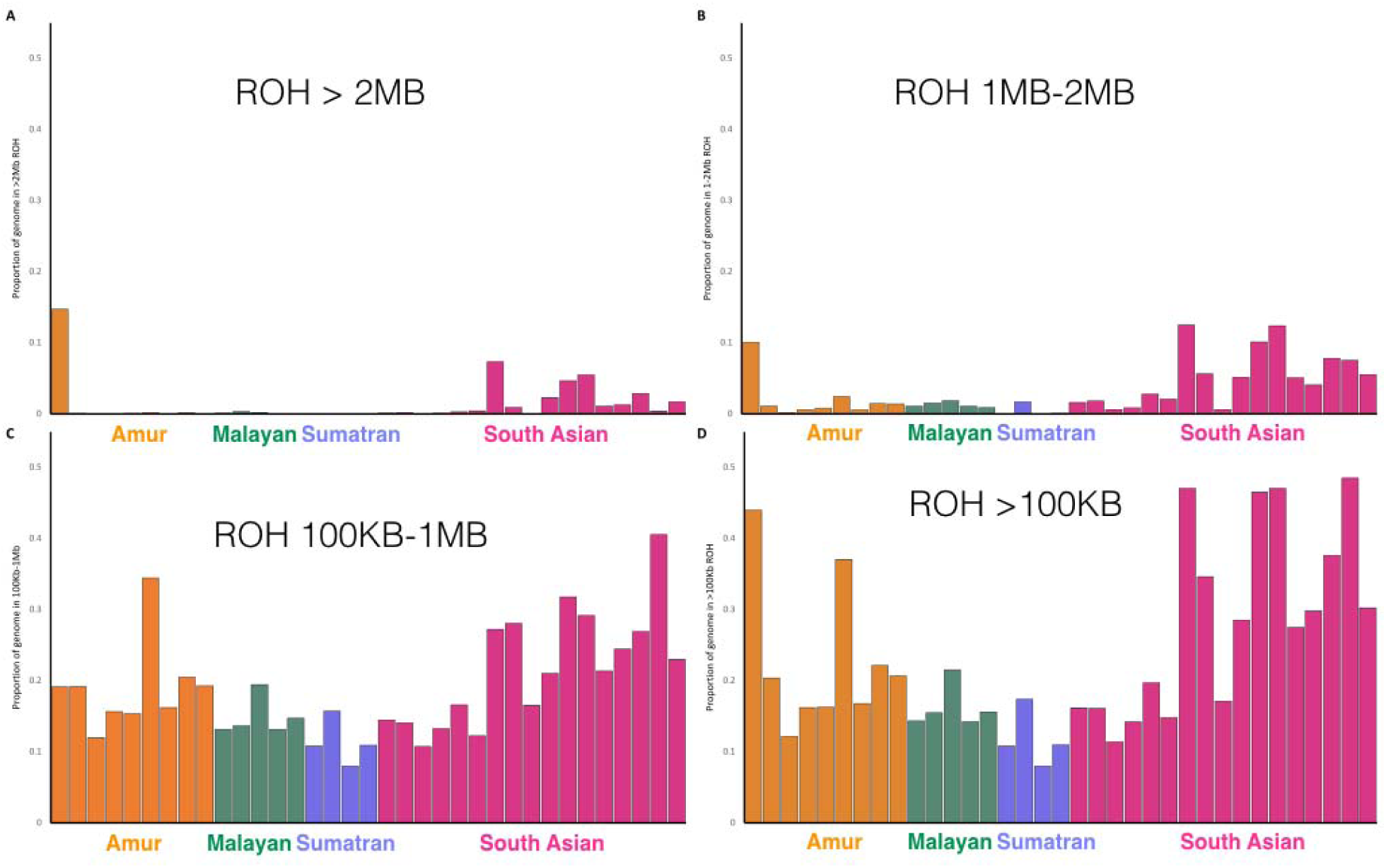
Runs of homozygosity inferred based on different run lengths. (A) > 2MB, (B) 1Mb-2MB, (C) 100KB-1MB and (D) Total ROH, which includes all run lengths greater than 100kb.

**Figure 3:**
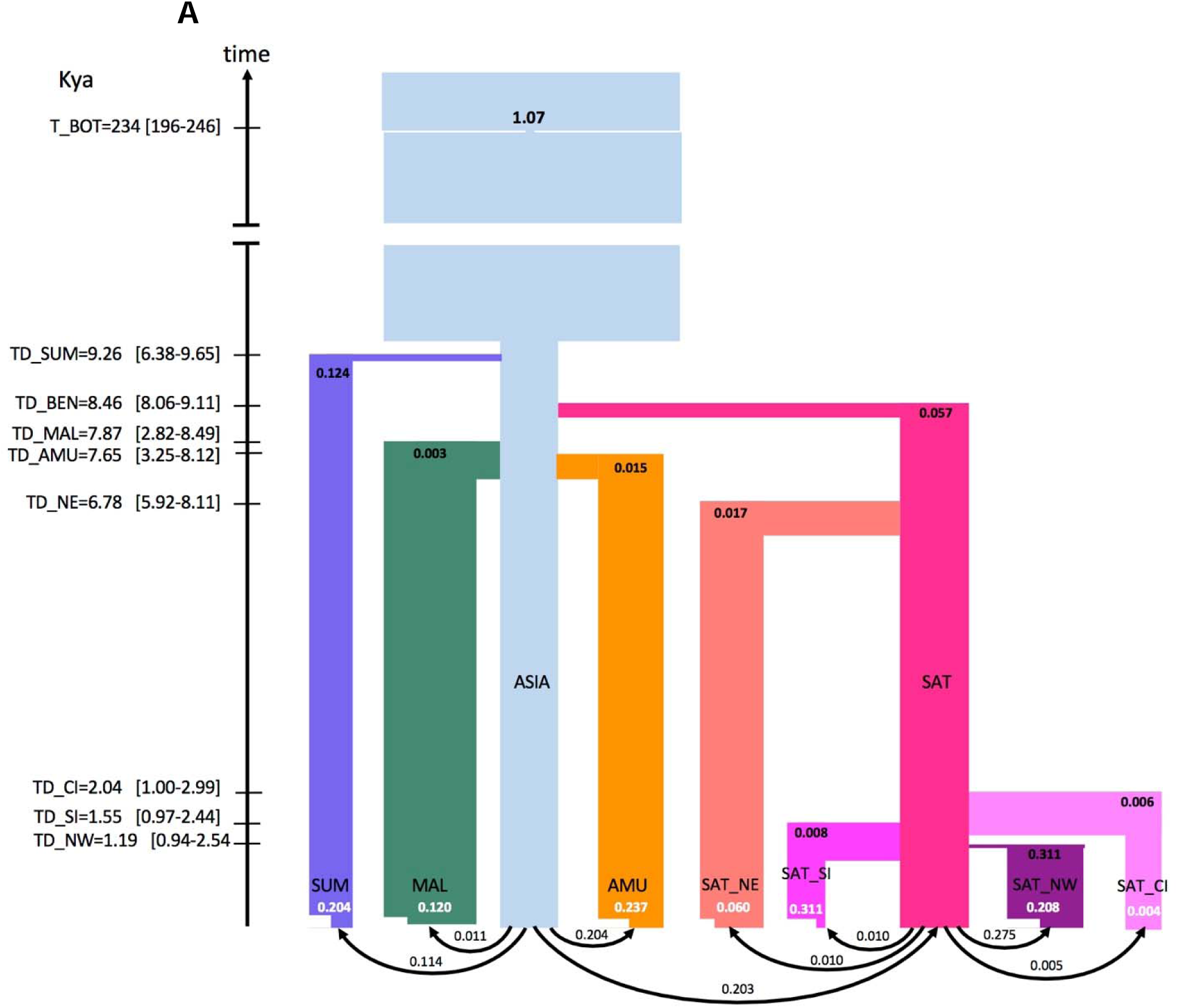
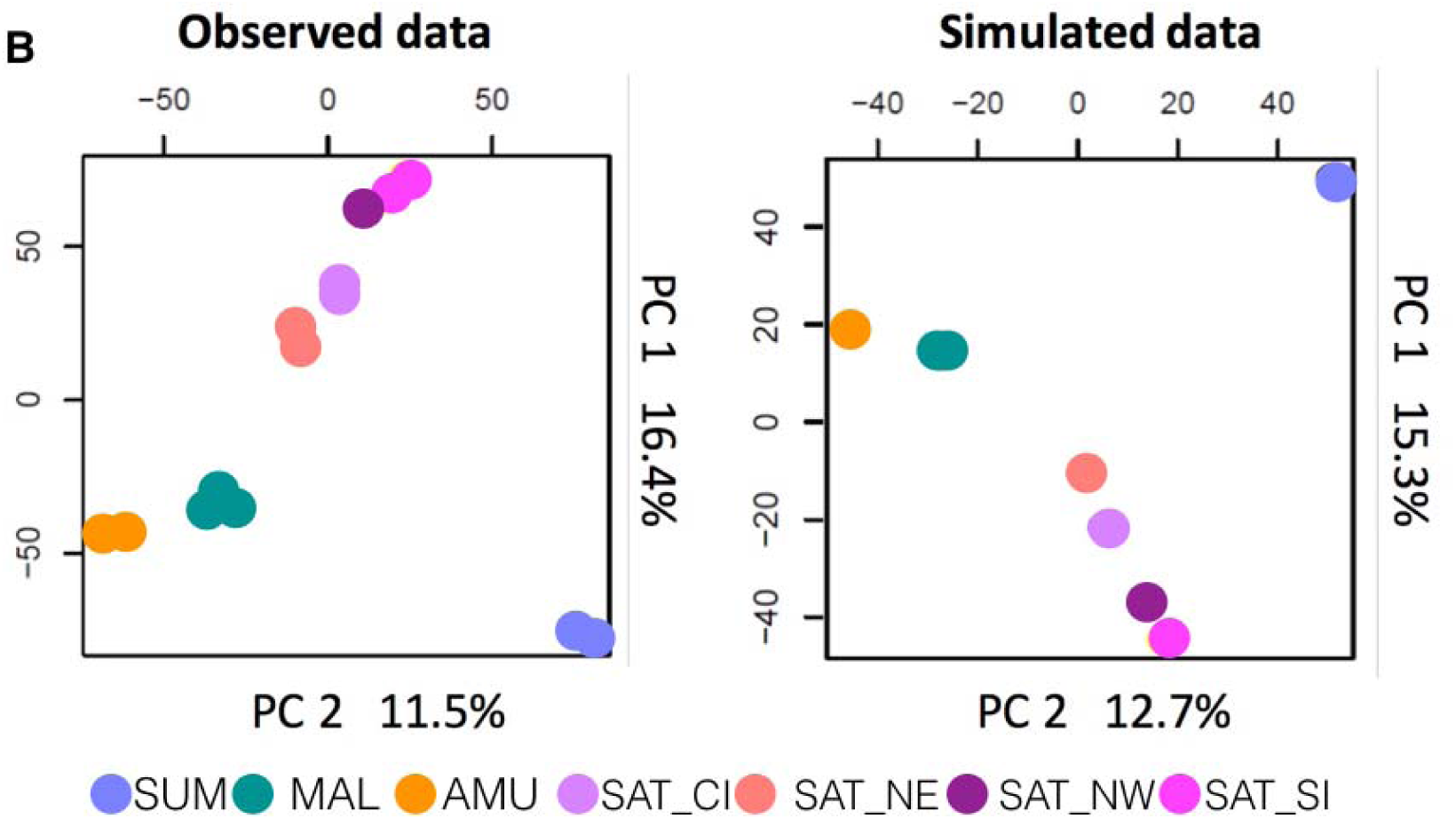
Estimated demographic history of Asian tigers: Sumatra (SUM: lavender), Malayan (MAL: dark green), Amur (AMU: orange) and ancestral South Asian (SA: hot pink), ancestral Asian metapopulation (Asia, light blue). The South Asian tigers further differentiated into North East (SA_NE, salmon pink), Central (SA_CI, light pink), Southern (SA_SI, dark pink) and North West (SA_NW, purple) populations. The approximate geographical locations of landscapes, modified from Natesh et al., 2017, shown in the inset map. (A) Founder effects are represented as horizontal lines with widths inversely proportional to intensity. Recent population contractions with intensity inversely proportional to current population size (t/2N) are reported in white text. Population bar widths are approximately proportional to estimated population sizes. Divergence (T_DIV) and bottleneck times (T_BOT) are reported in ky (thousand years ago), assuming a mutation rate of 0.35*10^−8^ and 5 years per generation. Times 95% CI values are shown within brackets on the left of the time arrow. Estimated values and associated 95% CI of all parameters are reported in Supplementary Tables 7 and 8 and (B) Comparison of PCA first 2 PC axes computed on observed and simulated data. The simulated scenario corresponds to that shown in Figure 3A, with parameter values taken from Supplementary tables 7 and 8.

PCA within subspecies (Supplementary Figure 5) suggested that South Asian tigers cluster into four sub-groups: (1) south India, (2) central and north India, (3) northeastern India, and (4) northwestern India. Some genomic sub-structuring was apparent in Malayan tigers, somewhat reflective of samples originating from the northern or southern Malayan peninsula (Supplementary Figures 5 and 6). Amur tigers did not demonstrate strong signatures of population sub-structuring (Supplementary Figures 5 and 6). Within subspecies, structure was confirmed in the additional PC axes for the full dataset (Supplementary Figure 4). While PC1, PC2 and PC3 separated putative subspecies (Amur, South Asian, Sumatran, and Malayan), PC1 & PC2 were also used in the supplemental figure 6B to separate the South Asian populations by geographic location clearly (northwestern India, south India and central, north, and northeast Indian tigers comprise three distinct groups).

Pairwise F_ST_’s (Supplementary Table 4) were approximately equal between subspecies and differences were consistent with geography. The F_ST_ between the Malayan and South Asian groups (0.164) was the lowest, while Amur and Sumatran F_ST_ (0.318) was highest, consistent with patterns seen in both ADMIXTURE and PCA. F_ST_ between putative South Asian tiger subpopulations in India (Supplementary Table 5) revealed high subdivision.

Since we sampled across landscapes within subspecies, we were able to compare population structuring within the four subspecies. Population genetic substructure is highest in the Indian subcontinent, while different geographical landscapes within other tiger subspecies are genetically the same (Amur) or less differentiated (Malayan). Our results contradict suggestions of population structure in wild Amur tigers (Sorokin et al., 2016), substantiate the significance of structure in South Asian tigers, and uncover hitherto unknown structure in tigers from the Malayan peninsula. Within South Asian tigers, we observed that northeast Indian tigers diverged considerably earlier than other South Asian populations. We reiterate that although these tigers are the most distinct of South Asian tigers, they are closer to South Asian tigers than they are to any other subspecies. The northeast Indian tigers in this study are from the state of Assam and sampling other, more eastern population from this remote region might yield interesting insights, as would samples from Indo-Chinese tigers.

### Genetic Variation and Runs of Homozygosity

We compared genome-wide variability between tiger subspecies/subpopulations to other cats (N=7) and endangered species (N=8, including endangered cats). Tigers had relatively high species-level genetic diversity (Supplementary Figure 7).

South Asian tigers had the highest nucleotide diversity (pi; Figure 1D), while Sumatran tigers had the lowest. Rarefaction analysis (ADZE; Szpiech et al. 2008) revealed that diversity estimates were approaching saturation for all populations (Supplementary Figure 12).

Historical demography and recent inbreeding are detectable through runs of homozygosity (ROH) in the genome (Pemberton et al 2012, Kardos et al. 2018). We quantified long (>2Mb) and shorter (100kb-1MB,1MB-2MB) homozygous stretches, as well as the proportion of more than 100Kb long ROH in the genome for several individuals (Figure 2). Somewhat surprisingly, individuals from the demographically large Indian tiger population revealed a high proportion of their genomes in long ROH, although variation in total ROH is high (Figure 2 and Supplementary Figure 8). Individuals from some populations (e.g. Central India) have low ROH, while some of the most inbred wild tigers in the world appear to be from India (e.g. from Ranthambore tiger reserve, Periyar tiger reserve and Kaziranga tiger reserve). Results were qualitatively similar when a sliding window approach was used to estimate ROH (Supplementary Figure 8).

Our data and analyses reveal South Asian tigers have the highest genetic variation across the genome. This is to be expected given historical records of South Asian tiger occupancy (Goodrich et al., 2015) across a large variety of habitats, where they subsist on a wide range of prey species that range from the large rhinoceros and gaur to the small hog deer and barking deer. Present population sizes of tigers in India and previous genetic studies based on a limited number of DNA microsatellite markers (Mondol et al., 2009) are also concordant with high genetic diversity in South Asian tigers. In contrast, certain South Asian tiger reveal signatures of potentially recent inbreeding, indicating population bottlenecks and isolation. High total genetic variation could be reflective of large numbers of tigers prior to intense hunting in India just a century ago (Rangarajan 2006).

A comparison among tiger populations is illustrative because it allowed us to reveal that Amur individuals do not harbor long homozygous stretches in their genomes, while individual South Asian tigers do. A closer look at landscapes and habitats in India and the Russian Far East reveal strong differences: India is dominated by variable habitats amidst a matrix of extremely high human population densities, while in the Russian Far East, human density is low and habitat is more continuous. Indeed, landscape genetics studies have suggested that high human population density is a barrier for tiger movement (Thatte et al., 2018). We suggest that extreme fragmentation and high human population density in India has resulted in isolated populations, where individuals may be more likely to mate with relatives. In contrast, despite low Amur tiger population densities in the Russian Far East, individual movement is not hindered by significant barriers and the population is more panmictic, with little to no sign of geographic population substructure.

Within South Asian tigers, we observe high variance in long ROH, generally thought to be the consequence of recent inbreeding, or possibly recent, intense bottlenecks (Kardos et al. 2018). For example, tigers from central India retain lower proportions of long ROH than those from other Indian landscapes (e.g. western India, southern India), possibly an outcome of higher recent connectivity between the central Indian tiger populations (Thatte et al. 2018, Yumnam et al. 2014), or lower historical bottleneck intensity. These very specific and hierarchical results underscore the importance of the inclusion of multiple genome-wide sampling across and within regions, as single representatives may be a poor reflection of inbreeding and variation for any given population, and do not provide a context with which to evaluate significance across subspecies and populations. In the future, simulations that incorporate realistic recombination rates could be used to model and disentangle the cumulative impacts of recent demographic history and very recent inbreeding on distributions of ROH in the genome.

### Demographic history of subspecies

We first reconstructed the past demographic history of each population with PSMC (Pairwise sequentially Markovian coalescent; Supplementary Figure 9), and our results paralleled those in Liu et al. (2018): all populations of tigers exhibit similar evolutionary patterns of population size decline.

Site frequency spectrum (SFS inferred from 259,499 SNP sites in non-transcribed regions at least 50 kb away from any known gene, selected to minimize the importance of background selection and GC-biased gene conversion) allowed us to investigate subspecies divergence, population size changes, as well as gene flow. The best fit scenarios contradicted previous results (i.e. Liu et al. (2018) suggesting divergence between subspecies around 68,000 years ago) by supporting a very recent (Holocene) divergence of all tiger subspecies (Figure 3A) from an ancestral population. Simulations supported a very strong bottleneck for the species occurring around 234,000 years ago, with most remaining lineages coalescing rapidly, consistent with a speciation event. This timing was consistent with signatures of population decline in the PSMC analysis (Supplementary Figure 9).

Existence of a large (theoretical) Asian metapopulation of tigers was followed by recent divergences between all four subspecies and between populations within the subspecies, including those within India. The best-fit scenario supported subspecies divergence in the Holocene between 7,500 and 9,200 years ago (i.e. 1,500 and 1,840 tiger generations). Sumatran tiger divergence correlates with sea levels rise (Heaney, 1991) and separation of the island of Sumatra. Note however that we imposed that this divergence post-dated the last-glacial maximum, i.e. 18,000 years ago or younger, because recent models of sea level rise suggest isolation from the mainland no later than 7,000 years ago (Bradley et al. 2016). Estimated migration rates were very low, with all populations receiving fewer than one migrant per generation; populations have been quite isolated since their initial early Holocene divergences. Additionally, we found that Sumatran and South Asian populations show evidence of a founding event, but Amur and Malayan populations do not. Both Sumatran and Amur tigers showed evidence of strong recent bottlenecks.

We further modeled the divergence within South Asian tigers into four populations: northwestern India, central India, southern India, and northeastern India. Because PCA suggests that central and north South Asian tigers are a single population, and north Indian tigers were not sequenced at high coverage, we left them out of these demographic analyses. We assessed the robustness of the northeastern population as being a part of the South Asian subspecies. In order to do so, the northeastern population was modelled as an independent subspecies, and allowed to diverge directly from the Asian metapopulation. However, such a model has a poorer fit to the data than if northeast Indian tigers are considered to be part of the South Asian subspecies (log_10_Likelihood difference between model is 37; Figure 3B). Within South Asian tigers, divergences are extremely recent (within the last 2,000 years), except for the northeast, which diverged early (6,800 years ago) after the separation of South Asian tigers 8,400 years ago from the ancestral Asian metapopulation. Within India, the northwestern population underwent a strong bottleneck at the time of its founding. Recent bottlenecks were most severe in the northwestern and southern populations, while the northeastern and central populations showed relatively weaker bottlenecks. Overall, tiger populations from all subspecies revealed signals of strong recent bottlenecks except central and northeastern South Asian tigers.

The variety of analyses we conducted (model-based inference, PCA, F_ST_, demographic modeling) revealed that tigers from different geographical locations are genetically distinct and have been isolated from each other for as much as 8,500 to less than 2,000 years (Figure 3A). Genomic divergences may reflect loss of connectivity due to sea level rise, which has separated the formerly continuous Sahul subcontinent of southeastern Asia into isolated islands, and changing environments due to human population size increase, including the rise of agriculture and climatic change of the mid-late Holocene.

Theoretical predictions (based on body size: Sutherland et al., 2000) and empirical results (genetics: Joshi et al., 2013; camera trapping: Singh et al., 2013) suggest that individual tigers can move extraordinary distances (e.g. 300 km), even across human-dominated landscapes. Such long-range movement would result in relatively low genetic differentiation if mating between members of separate populations was frequent and successful. However, despite the possibility of long-distance dispersal, our models suggest that migration rates between tiger populations have been relatively low, emphasizing separate recent evolutionary histories and that individual tiger movements may not represent the common population histories.

Although the timing and severity of the events differentiating tiger subspecies vary, our data and analyses confirm previous inferences (Liu et al. 2018) that the four tiger putative subspecies are valid both geographically and genetically, and that post-divergence gene flow has been relatively low. The order of divergence of the subspecies from the ancestral tiger metapopulation is partially consistent with previous suggestions of tigers being isolated in Sumatra first, likely due to sea level rise (consistent in sequence but not in timing with Liu et al., 2018), closely followed in time by those in India, then last by populations in Siberia and Malaysia (not consistent with Liu et al., 2018).

### Genome scans for selection

We investigated how genetic patterns might have been impacted by natural selection in the four tiger subspecies (Amur, South Asian, Malayan, and Sumatran). We computed a statistic, mPBS (metapopulation branch statistic, a simple extension of the PBS statistic of Yi et al. (2010), see Material and Methods), measuring the length of the branch leading to a given subspecies since its divergence from the others (Figure 4a and Material and Methods).

**Figure 4:**
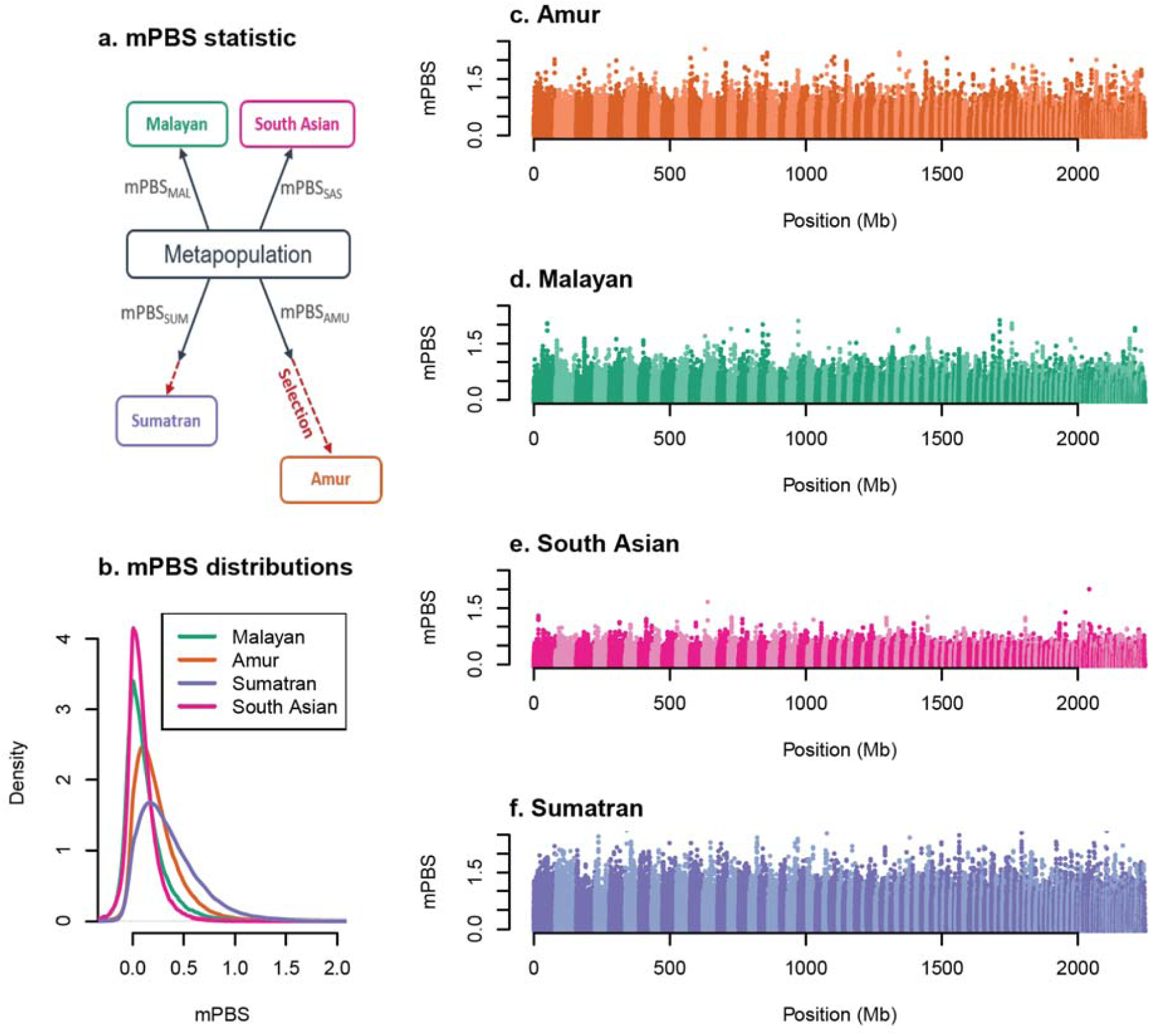
Genome scan for selection. (a) We present the mPBS statistic with a hypothetical model where the 4 populations diverge from a metapopulation, and where selection acts in both the Amur and Sumatra lineages, (b) the global distribution of observed mPBS for each population. Panels c to f correspond to the genome-wide distributions of the statistic for (c) Amur, (d) South Asian, (e) Malayan and (f) Sumatran tigers as a function of the genomic position. Alternating light and dark colors indicate different scaffolds.

The genome-wide distributions of the mPBS revealed that South Asian and Malayan populations had the lowest average values, suggesting short terminal branches subsequent to the divergence of these two populations from the hypothetical metapopulation (Figure 4b-f). On the contrary, Amur and Sumatran tigers had high values on average (Figure 4b-f).

We observed little difference between transcribed and non-transcribed regions in mPBS distributions, suggesting no strong differential impact of background or positive selection in tiger coding regions (Supplementary Figure 10). Both tails of the distribution are enriched (we did not filter for mutation types), possibly caused by biased gene conversion (Supplementary Figure 10). Note that average mPBS values were higher when considering only individuals with average coverage > 10X than when comparing fewer individuals with highest coverage (Supplementary Figure 10).

Overall, the mPBS distribution obtained under the neutral demographic model (Figure 4b) fit very well with the observed distribution (Supplementary Figure 11), implying that most observed differences between populations could be explained by their demographic history. We predicted high mPBS values in Amur tigers and Sumatran tigers where small effective sizes would yield high levels of genetic drift, but the observed values are even higher than those expected (Supplementary Figure 11), suggesting a possible effect of natural selection on genomic diversity in these subspecies. In contrast, we observed no apparent deviation of observed mPBS values from a purely neutral model in South Asian and Malayan populations.

Enrichment tests revealed an excess of moderately high values in Amur and Sumatran tigers rather than a few very extreme values, an observation that is compatible with the effect of polygenic selection rather than hard selective sweeps. In an attempt to identify biological functions putatively targeted by selection, we used functional enrichment tests (Daub et al. 2013, Gouy et al. 2017) based on mPBS values computed on all individuals (Amur and Sumatran) with average coverage greater than 10X. We mapped the top 0.1% regions with highest mPBS values to annotated genes (+/− 50 kb flanking regions). 119 and 80 genes (in Amur and Sumatran tigers, respectively) were found within these top 0.1% regions. We identified 15 statistically significant Gene Ontology (GO) terms in Amur tigers, and 5 in Sumatran tigers (Supplementary Table 9). Out of the 15 GO categories identified in Amur tigers, 4 have an unspecific function and the 11 others are involved in lipid processing and metabolism (Supplementary Table 9).

The genes responsible for the enrichment in fat metabolism-related GO terms were all included in the Cellular lipid metabolic process (GO:0044255). These included, for example, the Apolipoprotein B receptor (APOBR) or Caveolin-1 (CAV1) that are involved in the modulation of lipolysis. Fat metabolism enzymes included Phosphatidate phosphatase (LPIN2), Phospholipase B-like 1 (PLBD1), and Very-long-chain (3R)-3-hydroxyacyl-CoA dehydratase 2 (HACD2). We also identified genes involved in the mitochondrial respiratory chain: a Cytochrome P450 subunit (CYP1A2) and the mitochondrial Lipoyl synthase (LIAS). Cardiolipin synthase (CRLS1) is involved in the synthesis of cardiolipin, an important phospholipid of the mitochondrial membrane critical to mitochondrial function. Finally, Thromboxane-A synthase (TBXAS1) is involved in vasoconstriction and blood pressure regulation.

In Sumatran tigers, significant GO terms were related to cell development regulation: Regulation of neuron projection development (GO:0010975), Regulation of anatomical structure size (GO:0090066), and Regulation of cell development (GO:0060284). These four terms contain the same six genes: Tyrosine-protein kinase (RYK), E3 ubiquitin-protein ligase (RNF6), Low-density lipoprotein receptor-related protein 1 (LRP1), Angiotensin-converting enzyme (ACE), Rap1 GTPase-activating protein 2 (RAP1GAP2) and B2 bradykinin receptor (BDKRB2). These genes are involved in morphological development, and selection targeting these loci may help to explain why Sumatran tigers are in general smaller than other subspecies. Two other terms related to toxic substance processing, are: Response to toxic substance (GO:0009636) and Organophosphate biosynthetic process (GO:0090407).

### What evolutionary processes dominate the evolution of tigers and their subspecies?

Our models and analyses suggested relatively recent divergence between tiger populations (last 9,000 or so years versus 68,000 years inferred by Liu et al., 2018), highlighting the role of drift/stochastic processes in recent tiger evolution. Our inference is contingent on a mutation rate of 3.5 × 10^−8^ from Liu et al., 2018. Discrepancy between our and Liu et al., 2018 estimates could also be due to differences in filtering criteria (Liu et al., 2018 use min DP=4, min GQ=20, whereas we have used minDP=10, and min GQ=30) or the sites considered for the analyses. We used a restricted set of sites that were far from coding regions, and thus minimally affected by background selection and biased gene conversion (about 100Mb worth of data), whereas Liu et al. 2018 used about 44 Mb worth of data (for GPhos analyses) without control for genic regions, background selection or biased gene conversion. Our results consistently underline the genome-wide importance of genetic drift. Despite recent divergence we found significant genetic differentiation between tiger populations, possibly because of the intense bottlenecks these populations have experienced.

Our results suggested that Amur tiger genomes demonstrate signals of selection, with possible adaptations to colder environments. Genes and pathways involved in lipid metabolism are under selection in two human populations that live in cold environments, including Greenlandic Inuit (Fumagalli et al., 2015) and Indigenous Siberians (Hallmark et al., 2018), which revealed signatures of selection on. Polar bears genomes also reveal signatures of selection on lipid metabolism genes (Liu et al., 2014). Understanding the distribution of adaptive variants could be important for future conservation efforts, especially if priority were placed on preserving these cold-adapted populations, which may be disadvantaged under future warming scenarios

Sumatran tigers appear to have experienced strong genetic drift following vicariance from mainland southeast Asia, maintained a smaller effective population size, and have experienced a strong recent bottleneck. Although Liu et al., (2018) suggested that selection for body size targeted the *ADH7* gene, we did not detect any signature of selection at this locus in our Sumatran samples. However, we identify alternative candidate genes that are potentially involved in body size, such as the genes found in the *Regulation of anatomical structure size* GO term (Supplementary Table 9). We caution that it is difficult to truly distinguish among all population genetic processes, especially selection, without more data, and assignments of GO categories designed from model organisms are only a substitute for more definitive tests of selection. Differences between our study and Liu et al. (2018) may be due to the improved quality of the genome we built and mapped to, which generally increases the accuracy of gene finding and annotation software.

We did not detect signatures of selection or extensive gene flow into Malayan and South Asian tiger genomes, suggesting that their genomic variation was due primarily to drift. Indian tigers appear to have experienced intense founding events, intense recent bottlenecks, and population structuring, suggesting a relatively stronger role for drift (compared with Malayan tigers) in shaping genome-wide variation.

### Conservation implications

We show that tigers have recently differentiated into four subspecies with unique gene pools created by contrasting histories of drift and selection that has made each subspecies evolutionarily unique. Our results suggest common and varied conservation strategies for Amur, Malayan, South Asian (Indian) and Sumatran tigers might be considered. For Amur tigers, our results from population structure analyses, demographic history based divergence and signatures of possible selection confirm (as suggested by Liu et al. 2018 and Wilting et al. 2015) their separate management status. Increasing population size and enabling gene flow over the long term might augment the currently low genetic diversity in this population. In contrast, restoring and maintaining gene flow between populations through habitat corridors may be more important along with increasing population numbers for South Asian tigers, where genetic variation is high, but connectivity is low and runs of homozygosity suggest recent inbreeding. Recent fragmentation and ensuing loss of connectivity appears to result in significant autozygosity in the South Asian tiger, and assisted geneflow could be considered as a management strategy, especially when inbreeding is associated with loss of fitness and potentially inbreeding depression. Within South Asian tigers, we suggest that the management status of northeast Indian tigers be re-evaluated given their antiquity and potential genetic distinctiveness (Kolipakam et al. 2019). The surprisingly high (relative) genetic variation and population differentiation in Malayan tigers bodes well for their future survival. It will be critical for future conservation efforts to prioritize population recovery and gene flow through connectivity, and to ensure increased population sizes. Critical to such action is a better understanding of within population genetic variation using spatially-explicit, non-invasive sampling. Finally, Sumatran tigers should be managed separately because like Liu et al (2018) and Wilting et al (2015), our results re-iterate their uniqueness. Their genomes show signatures of selection for genes regulating body size (broadly consistent with the findings in Liu et al (2018)).

Our analyses also suggest that introgression from other gene pools into Amur and Sumatran tigers may result in outbreeding depression and/or loss of their unique adaptations. Poleward progression of subtropical climates may favor adaptive alleles (found in more southern Sumatran tigers) in more northern populations (e.g. Amur, Malayan or Thai tigers), potentially increasing the value of these adaptations to future survival. Ongoing human impacts like fragmentation will likely continue to disrupt natural evolutionary processes. Managing local populations to minimize human impacts that allow continued tiger evolution may be the key to species survival, and the important conservation strategy for the Anthropocene.

## Materials and Methods

### Sample Collection

We obtained tissue, blood, or serum samples from as many geographically distinct tiger populations as possible. This amounted to 65 samples from 4 tiger subspecies including 21 Indian South Asian tigers (*P. t. tigris*), 19 Eastern Siberian tigers (*P. t. altaica*), 15 Malayan tigers (*P. t. jacksoni*), and 11 Sumatran tigers (*P. t. sumatrae*). A list of final samples sequenced and their sources are available in Supplementary Table 1. We also included 1 already sequenced sample, which brought the sample total to 66 (see Supplementary Table 1).

### Reference assembly sequencing and de novo assembly

In order to better understand genome-wide variation and call variants reliably, we first built a new tiger genome assembly using the 10X Genomics Chromium Platform for a wild-caught Malayan individual. We received whole blood from a wild born Malayan tiger (*P. t. jacksoni*) sampled by the El Paso zoo, Texas on 7/28/2016, collected as part of a routine health checkup. We immediately froze the sample at −80°C until it was shipped on dry ice to the Barsh lab at HudsonAlpha for extraction and delivery to the Genome Services Lab (GSL) at HudsonAlpha Institute for Biotechnology, Huntsville, Alabama. DNA was extracted and purified using the Qiagen MagAttract HMW DNA kit. GSL staff prepared a linked-read sequencing library using the Chromium controller. The library was sequenced on one lane of a HiSeqX. We assembled the genome using the SuperNova assembly software (1.1.4) provided by 10x Genomics using the standard pipeline. We refer to this assembly as Maltig1.0 hereafter.

Based on Assemblathon2 statistics (Bradnam et al. 2013), this improved assembly corresponded to a 3.5-fold increase in the contig N50 value to 1.8 Mb and a 2.5-fold increase in the scaffold N50 value to 21.3 Mb (as compared to Cho et al. 2013; Supplementary Table 2). In addition, the resulting assembly had ∼1% fewer ambiguous bases across all scaffolds (Supplementary Table 2). Details of samples used for various analyses are in Supplementary Table 3.

### Whole Genome Re-sequencing and Variant Discovery

Details on DNA extraction, library preparation, and variant discovery methods can be found in supplementary materials.

### Population structure

We first investigated admixture and structure between populations using Plink2 (Chang et al. 2015). We used VCFtools to filter the initial variant call file using ‘max-missing 0.95’ and ‘maf 0.025’ to remove sites with missing data and rare variant calls. We then converted to Plink’s ‘.ped/.map’ format using VCFtools, and subsequently converted to ‘.bed/.bim/.fam’ format within Plink2 using the flag ‘--make-bed’. PCA was then run on the resulting bed file using the flag ‘--pca 10’ which computes the variance-standardized relationship matrix. PCAs were then plotted using R. For smaller runs an additional step was added within Plink2 to first calculate the frequencies using the flag ‘--freq’. Subsequently, PCA was run using the ‘--pca’ flag and inputting the frequency file using the ‘--read-freq’ flag. We used this protocol on the vcf with all individuals and subsequently, we divided the vcf into the putative subspecies for within subspecies runs.

The program ADMIXTURE was used to infer structure between populations and inform clusters which represent populations with distinct histories (Alexander et al. 2009). ADMIXTURE uses maximum likelihood-based models to infer underlying ancestry for unrelated individuals. We used the filtered dataset (VCFtools max-missingness cutoff of 95%, minor allele frequency cutoff of 0.025) and resulting Plink formatted files for input into the software. In order to infer the most likely value of K, values of 2-8 were run. We also performed K validation in order to compute the cross-validation error for each value of K, by using the ‘--cv’ flag within the program. The value with the least error is informative of the best value of K for the data.

### Rarefaction analysis

To ensure that our data was reflective of the diversity within each subspecies/unit as defined by ADMIXTURE, we used the program ADZE (Szpiech et al. 2008). ADZE runs a rarefaction analyses on polymorphism data in order to estimate the number of alleles private to any given population (not found in any other population), considering equal-sized subsamples from each input population. In addition, the program calculates distinct alleles within each population. We calculated the private alleles across the four main populations/sub-species as designated by the ADMXITURE software, in addition to the distinct alleles within each of the four populations individually.

### Population differentiation and diversity

We calculated pairwise F_ST_ between each subspecies group as defined by ADMIXTURE using VCFtools. Variant call data was subdivided into sub-species based on PCA (South Asian, Sumatran, Amur, and Malayan as subgroups) and was used to compute pairwise F_ST_ between each group. Nucleotide diversity (pi) was calculated using VCFtools.

In order to detect the number of single nucleotide variants (SNV), the data were filtered using VCFtools (Danecek et al. 2011) to a minimum base quality of 30, genotype quality of 30 and depth 10. We additionally filtered for minor allele frequency of 0.025 and allowed a maximum 5% missing data in any loci. RTG tools (https://www.realtimegenomics.com/products/rtg-tools) vcfstats was used to calculate the total number of heterozygous SNP sites for each individual. These values were then plotted alongside comparable estimates for other species reported in Abascal et al. (2016).

### Ancient demographic history

Pairwise sequentially Markovian Coalescent (PSMC) (Li and Durbin, 2011) is a single genome method to detect historical effective population size. In order to estimate historical population size changes for the different subspecies, we removed sex chromosome scaffolds for AMU1, MAL1, SUM2 and SAT_SI3 (the highest coverage individual for each subspecies). The procedures for sex chromosome filtering can be found in the supplementary text (and Supplementary Figure 1). Additionally, sites with a minimum of half the average sequencing depth or twice the average sequencing depth were filtered out while calling variant sites. The resulting scaffolds were then used to estimate the effective population size across 34 time intervals as described in Li and Durbin (2011). 100 rounds of bootstrap replicates were performed.

## Runs of Homozygosity

To estimate runs of homozygosity (ROH), we used the filtered SNPs from the autosomal scaffolds. Individuals with more than 10x average coverage were grouped as per subspecies. We used BCFtools/RoH (Narasimhan et al. 2016) to estimate ROH. The autozygous runs obtained were classified into various lengths (runs between 10kb and 100kb, runs between 100kb and 1 Mb, and runs longer than 1 Mb). Proportion of genome in ROH for 1Mb was estimated as the total length of the genome in more than 1Mb runs divided by the total length of autosomal scaffolds. Similar calculations were made for 100kb to 1Mb runs, and for 10kb-100kb runs except the length of the genome longer than 1Mb and 100Kb were subtracted from total length of autosomes respectively. We used an additional sliding window approach, details of which can be found in the supplementary methods.

## Demographic history with SFS and coalescent models

### Demographic models

Data filtering procedures for the demographic models can be found in the Supplementary text. Using the program fastsimcoal 2 (Excoffier et al. 2013), we performed demographic estimations of the model shown in Figure 5 on two datasets in two consecutive steps, such as to reduce the number of parameters to estimate simultaneously. The first step consisted in estimating the demography (24 parameters) of four tiger subspecies (Malaysia – MAL, Sumatra –SUM, South Asian – SAT, and Amur – AMU) using the individuals of each subspecies that had the highest coverage. We thus selected 3 SUM individuals, 3 SAT individuals from South India (SAT_SI), four MAL individuals, and 3 AMU individuals, which all had >20X coverage on average (See list in Supplementary Table 3). We modeled the four-subspecies as belonging to a large Asian metapopulation, from which they would have diverged some time ago while still receiving some continuous gene flow from the metapopulation. Note that this continent-island population structure amount to modeling a set of populations having gone through a range expansion (Excoffier 2004). We assumed that each of the four subspecies could have gone through two distinct bottlenecks, one that would have occurred at the time of the separation from the Asian metapopulation, to mimic some initial founder effect, and one that would be recent to mimic habitat deterioration. We also assumed that the Asian metapopulation could have gone through an ancestral bottleneck sometime in the past.

The second step used estimated parameters in a more complex model including the specific demography of four South Asian tiger populations (24 new additional parameters). To this aim, as in the previous analysis, we selected individuals with the highest coverage (>20X) from each population (see Supplementary Table 1, samples used represented in Supplementary Table 3). No individuals from SA_NOR were included as their coverage was low and they are part of the same genetic cluster as SA_CI. In order to correctly estimate the relationship between these populations and the other subspecies, we also included 3 MAL individuals in this analysis. The new model included all the parameters from the previous model, fixed at their previously estimated values, except some parameters re-estimated for the SA_SI population, which was now assumed to have diverged from an Indian metapopulation at some time in the past, like the other three SA tiger populations. We also estimated the size and the divergence of the SA metapopulation from the Asian metapopulation. We allowed the sampled SA populations to have gone through two bottlenecks (an initial founder effect and a recent bottleneck). The parameter estimated in these two steps are shown in Supplementary Tables 7 and 8) and the resulting demography is sketched in Figure 3.

### Parameter estimation and fastsimcoal2 command line

*Fastsimcoal2* (Excoffier et al. 2013) was used to estimate parameters from the multidimensional site frequency spectrum (SFS) computed on non-coding regions at least 50 Kb away from known genes. The multidimensional SFS was computed with the program Arlequin ver 3.5.2.2 (http://cmpg.unibe.ch/software/arlequin35) on polymorphic sites matching the filtering criteria listed above. In order to infer absolute values of the parameters, we used a mutation rate of 0.35e-8 per site per generation estimated in a previous paper on tiger demography (Liu et al. 2018), and we assumed that the proportion of monomorphic sites passing our filtering criteria in each 1Mb segment was identical to the fraction observed in polymorphic sites. Parameter estimation was obtained by maximum-likelihood estimation obtained from 100 independent runs of *fastsimcoal2*, 60 expectation conditional maximization (ECM) cycles per run and 500 thousand coalescent simulations per estimation of expected SFS. The *fastsimcoal2* command line used for the estimation was of the type fsc -t xxx.tpl -n500000 -d -e xxx.est -M -l30 -L60 -q -C1 --multiSFS -c1 -B1 where fsc is the *fastsimcoal2* program and xxx the generic name of the input files. The.est and .tpl input files used for inference on the two datasets are listed in the Supplementary material.

Confidence intervals were estimated via a block bootstrap approach. We generated 100 bootstrapped SFS by resampling (and adding) the SFS from 1 Mb segments of concatenated non-coding segments along the genome, and for each of these bootstrapped samples we re-estimated the parameters of the model using 10 *fastsimcoal2* independent runs starting at the maximum-likelihood parameter values. Again, we used 60 ECM cycles for each run and performed 500,000 simulations for estimating the expected SFS under a given set of parameter values. The limits of 95% confidence intervals were estimated by computing the 2.5% and 97.5% quantiles of the distribution of 100 maximum-likelihood parameter values.

### Genome scan for selection

To detect the footprints of natural selection in different tiger subspecies, we analyzed individuals with coverage > 10X from 4 subspecies (n = 34). We filtered out genotypes with depth of coverage < 10 (DP) and genotype quality < 30 (GQ). We excluded scaffolds shorter than 1 Mb. We kept sites with no missing data among the 34 individuals.

We considered the 4 subspecies as 4 populations and computed pairwise F_ST_ values along the genome over 50 kb sliding windows (with a step of 10 kb) using the R package PopGenome (Pfeifer et al. 2014). We then computed a measure of selection similar to the Population Branch Statistic (PBS) (Yi et al. 2010). The PBS statistic is based on a three-population comparison and measures the length of the branch leading to a given population since its divergence from the two other populations. This statistic is not able to accommodate more than three populations and relies on a tree-based model that does not correspond to tigers’ demographic history. Therefore, we extended this statistic to the case of four populations under a more suitable model than a tree-based one. Furthermore, using all four populations allows to better characterize the differences that are exclusive to specific branch. We define:

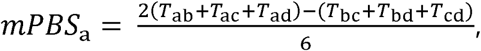

where *T*_*ij*_ is the divergence time, in generations, between population *i* and *j* (Nei, 1972):

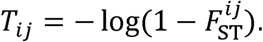

This statistic assumes that each population diverged from a metapopulation independently and that no migration occurred following divergence. It measures the length of the branch leading to a given lineage since its divergence from the metapopulation. Selection in a given lineage will lead to a much longer terminal branch than under neutrality. This would translate to extreme mPBS values.

To compare observed mPBS values to expectations under the tigers’ demographic history, we simulated 1 million genomic windows using the demographic model inferred previously. Window size and sample sizes for each population are the same as in the observed dataset. Parameter values are fixed and correspond to the maximum likelihood estimates (Supplementary Tables 7 and 8). Then, we computed the mPBS statistic for each population to generate a null distribution. Observed and simulated distributions were then represented to see whether observed values deviated from neutral expectations.

Enrichment tests were then used to detect the targets of selection. These tests are a conservative approach to detect selection because they are less susceptible to the influence of non-selective forces. To identify putative genes under selection, we considered predicted genic regions of the tiger genome for which a homolog has been annotated using Exonerate (protein2genome model). To avoid spurious enrichment signals due to the presence of multiple homologs for a single gene, we kept only one homolog for each predicted gene. If different homologs on the same strand overlap, we pick the first one and ignore the others. We retained a total of 12,771 genes after filtering.

We also checked whether some Gene Ontology (GO) terms (Ashburner et al. 2000, Mi et al. 2016) were enriched across candidate genes (Fisher’s exact test performed on human GO terms). Genes (+/− 50 kilobases flanking regions) were considered as candidates if they overlapped with a window that was in the top 0.1 % of mPBS value of a given population. The reference list of genes for the enrichment test is set as the list of genes after filtering (12,771 genes).

## Acknowledgements

We thank the American Zoo Association for endorsing our research and collection of samples from captive tigers, Tara Harris (then Minnesota Zoo) and Kathy Traylor-Holzer (Tiger Species survival program) for help with captive tiger studbooks. We thank San Francisco Zoo, San Diego Zoo (BR2016035; Leona Chemnick for assistance with DNA extraction and sample transport), El Paso zoo, Omaha Zoological Society, WCS Bronx zoo (IC2016-0464 WCS; Dee McAloose and Jean Pare for assistance with sample transport), Gopala Battu for assistance with sequencing and sample preparation at Hudson Alpha. Zachary Szpiech for assistance with ADZE. Infosys Travel Award to A Khan, SciGenome Research Foundation Grant to A Khan, Fulbright Nehru Academic exchange fellowship to U Ramakrishnan, CEHG, Stanford University funding to U Ramakrishnan, Wellcome Trust-DBT Indian Alliance Senior fellowship to U Ramakrishnan (IA/S/16/2/502714), Genomics Facility of CCAMP. We thank Atul Upadhayay for bioinformatics support, and the computing facility at NCBS.

## Author contributions

EEA, AK, RWT, UR, and EAH designed the study. EEA, AK, RWT, AG, GG, AT, JTLK, SR, SP, GB, CK, SP, AC, LE conducted lab work and analyses. GB, CK, MG, DM, AZ, UB, AR, EL, OAR, YVJ, EAH, UR provided samples and funding. EEA, AK, RWT, AG, GG, AT, JTLK, SR, SP, GB, CK, SP, AC, MG, DM, AZ, UB, AR, EL, OAR, YVJ, DP, LE, EAH, UR wrote and edited the paper.

