## Supplementary material for "Recent evolutionary history of tigers highlights contrasting roles of genetic drift and selection"

**Supplementary methods**

**DNA Extraction and Library Preparation**

We extracted DNA from samples using the Qiagen DNeasy blood and tissue kits (Catalogue #69504) and quantified DNA concentrations with the Qubit dsDNA HS assay kit (Q32851). As a number of our samples yielded very low amounts of DNA (< 1 ng), we used an approach that scales down the input reagents and necessary DNA input and modified the Illumina Nextera kit protocol (Baym et al. 2015). Genomic libraries were run on a 2100 BioAnalyzer (Agilent Technologies using High Sensitivity DNA Chips (Catalog #5067-4626) to determine the quality, quantity, and fragment size distributions. Libraries were sequenced on Illumina HiSeqX, HiSeq 4000, and HiSeq 2500 for between 5-25x coverage using paired-end 2x150bp reads (Supplementary Table 1).

**Variant Discovery**

We trimmed reads prior to mapping with TrimGalore (Krueger 2015), then mapped reads to the Maltig1.0 reference genome using BWA-MEM (Li 2013) and sorted and indexed using SAMtools (Li et al. 2009). We marked duplicate reads with the Picard Tools `MarkDuplicates` command (<http://broadinstitute.github.io/picard>). We then called variants from the resulting BAM files using FreeBayes (Garrison & Marth 2012). The resulting VCF file was filtered with VCFtools (Danecek et al. 2011) to a minimum quality of 30, a genotype quality of 30, maximum of 2 alleles, Hardy-Weingburg p-value of 0.0001, minimum allele frequency of 0.01, and a minimum minor allele count of 3. The pipeline was managed and parallelized using NextFlow (Tommaso et al. 2017).

**Sex chromosome filtering for PSMC analysis**

We excluded the sex chromosomes for demographic analyses because their effective population size differs from that of autosomes. We identified potential sex chromosome scaffolds using three strategies: (a) The assembled tiger scaffolds were aligned to the domestic cat genome assembly (Felis_catus_9.0) using BLAST (Altschul et al. 1990). The top hits for each alignment were chosen. All tiger scaffolds that aligned with the domestic cat X chromosome scaffolds were marked as a sex chromosome; (b) The re-sequencing reads for MAL1 (also the individual used for 10x assembly) were aligned to the tiger genome assembly. All the scaffolds sequenced at half the average depth or twice the average depth were marked as sex chromosome scaffolds; (c) Since the sex for many tigers in this dataset are known, a statistical association between scaffolds at half the average sequencing depth and zero sequencing depth was established (Supplementary method Figure 1). Females had few scaffolds sequenced at zero depth, and these represent Y chromosomes. Males had certain scaffolds sequenced at half the average depth, representing potential X chromosome. Using a conservative approach, all the scaffolds identified in even one of these analyses were marked as sex chromosomes and removed.

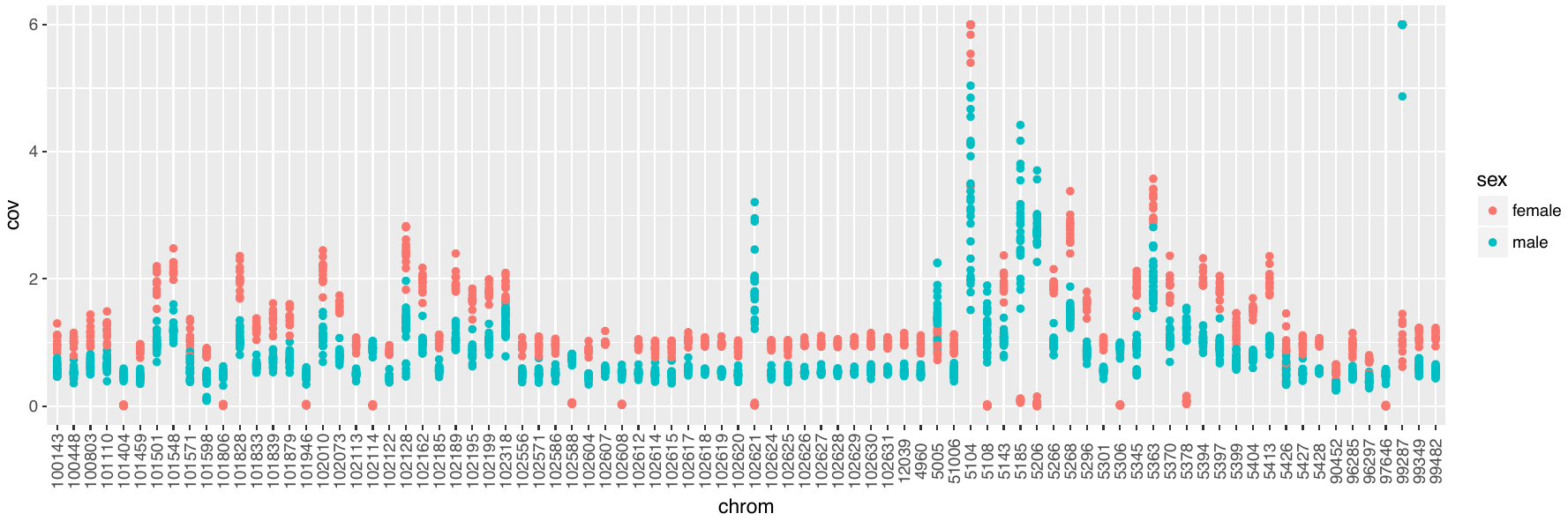

**Supplementary Figure 1**: Association between sex and coverage of various scaffolds. Coverage is normalized per individual.

**Estimation of ROH using sliding window approach**

Within individual genomes, we identified ROH using the methods first implemented in Pemberton et al. (2012). For each of the four tiger populations, we estimated the allele frequencies at each SNP using the observed allele frequencies for the four populations in our data set. To identify ROH, we employed a likelihood method adapted by Pemberton et al. (2012). This approach, which forms the basis of the ROH inference program GARLIC (Szpiech et al. 2017), considers a sliding window of *n* SNPs that moves along the chromosome in single SNP increments. To ensure robustness in our results, we repeated the ROH identification procedure with three values of *n*: 100, 250, and 700. Within each window, a log-likelihood score was computed for each SNP, comparing the hypothesis that the SNP is autozygous to the hypothesis that it is non-autozygous, allowing for an error term that accounts for mutation or genotyping error. As in Pemberton et al. (2012), we set the error parameter to 0.001. The overall score of a window is then the sum of the scores of the SNPs it contains, with an observed homozygous SNP contributing a positive score (unless the SNP is monoallelic in the population in question), and an observed heterozygous SNP contributing a negative score.

Following that, all windows with an overall score of 0 or greater were taken to be in an ROH, with overlapping windows merged and considered as part of a single ROH. The bp length of each ROH is taken to be the length of the interval between the its two most extreme SNPs, including the endpoints. For the three values of *n*, we then plotted the total length of ROH present in each individual, as well as the total length of long ROH (≥ 1Mb) present in each individual (Supplementary Figure 8).

**Demographic history with SFS and coalescent models**

*Data filtering*

In order to minimize the effect of background selection (BGS) that could bias demographic inferences (Pouyet et al. 2018), we excluded genic regions and those within 50kb of gene transcripts. We concatenated the remaining sequences in a single fictive chromosome and divided it into smaller segments of 1Mb in length. These 1Mb segments were used for block-bootstrap resampling for the estimation of parameter confidence intervals (see below).

From the VCF including information on polymorphic sites, we used the Snow leopard genomic sequence (Cho et al. 2013) as a reference to filter out triallelic sites and those sites for which the Snow leopard variant was missing. We further excluded sites where any individual had a site with a coverage (DP) lower than 10 or a genotype quality (GQ) lower than 30. To exclude the effect of GC-Biased Gene Conversion (BGC) on demographic inference, (Pouyet et al. 2018), we only considered sites with A ↔ T or C ↔ G mutations that are not affected by BGC. Finally, for any given data set defined below, sites with any missing data for one of the filtering step described above were also excluded. The number of monomorphic sites passing all our filtering criteria including BGC was computed by assuming that BGC-free monomorphic sites would have a similar frequency among all monomorphic sites than BGC-free polymorphic sites among all polymorphic sites.

**Other supplementary figures**

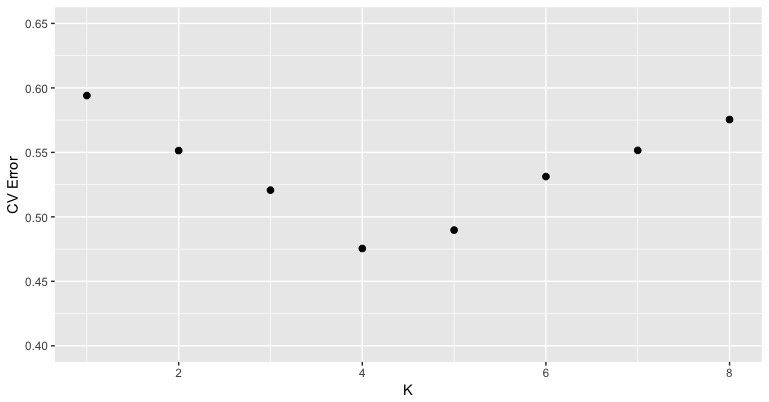

**Supplementary Figure 2**: Cross-validation error for runs of ADMIXTURE across different values of K.

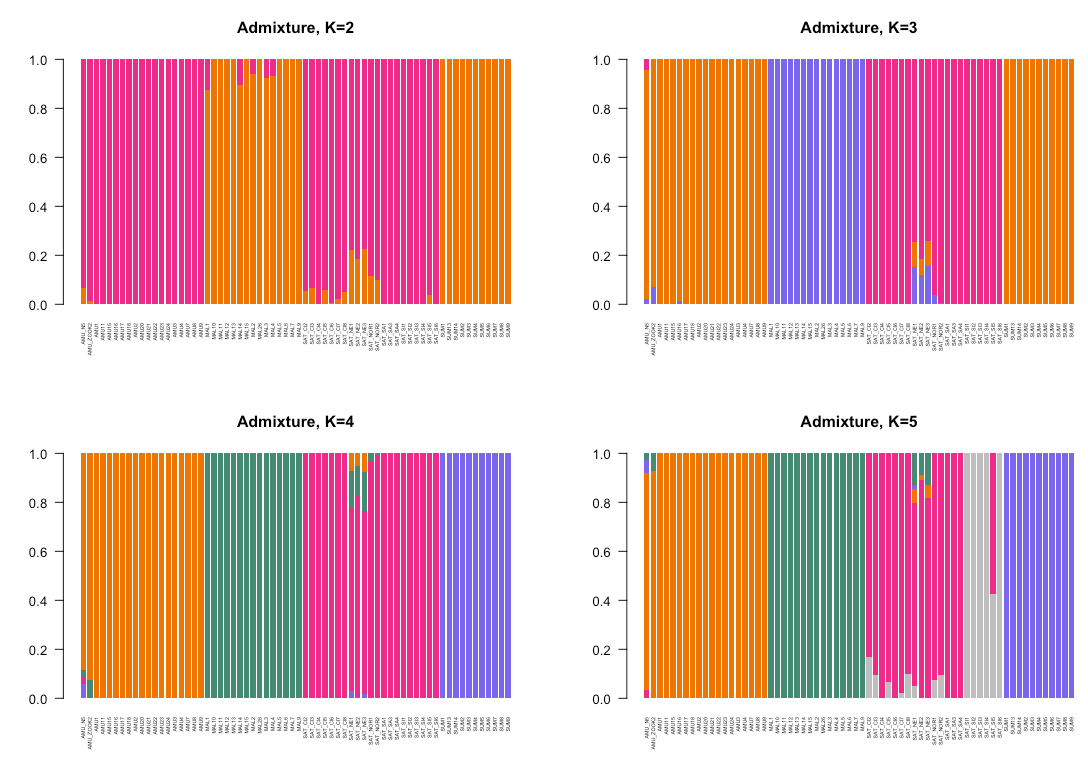

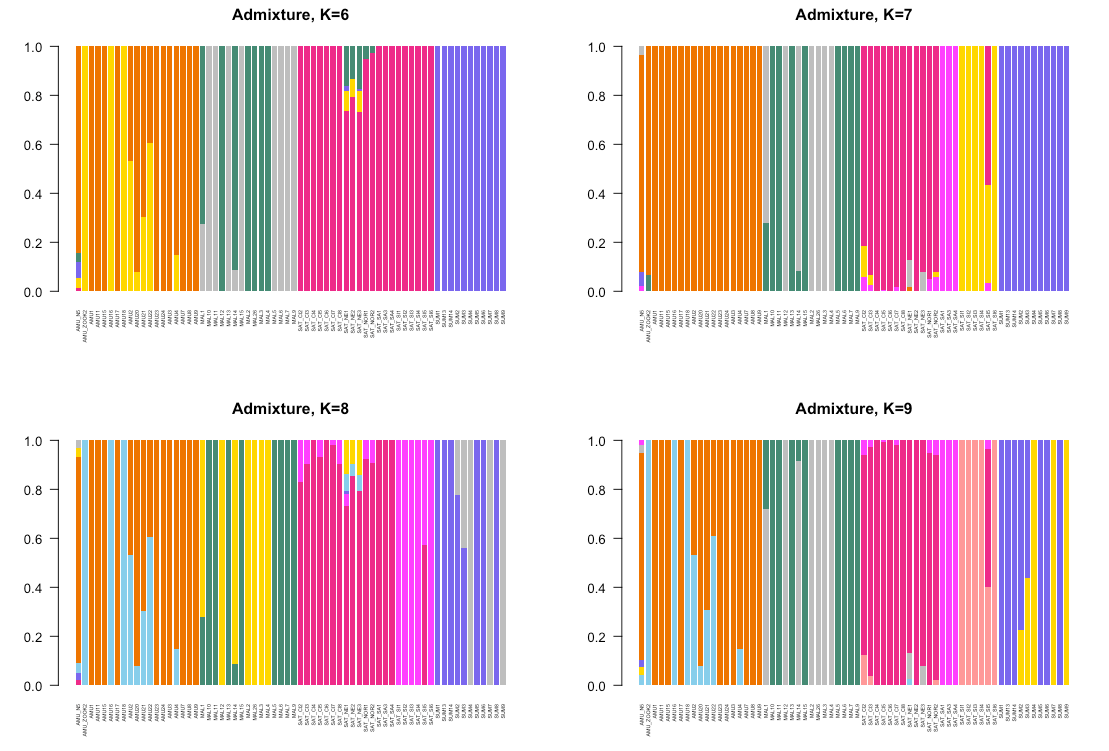

**Supplementary Figure 3**: ADMIXTURE plots for all tigers for K values 2-9.

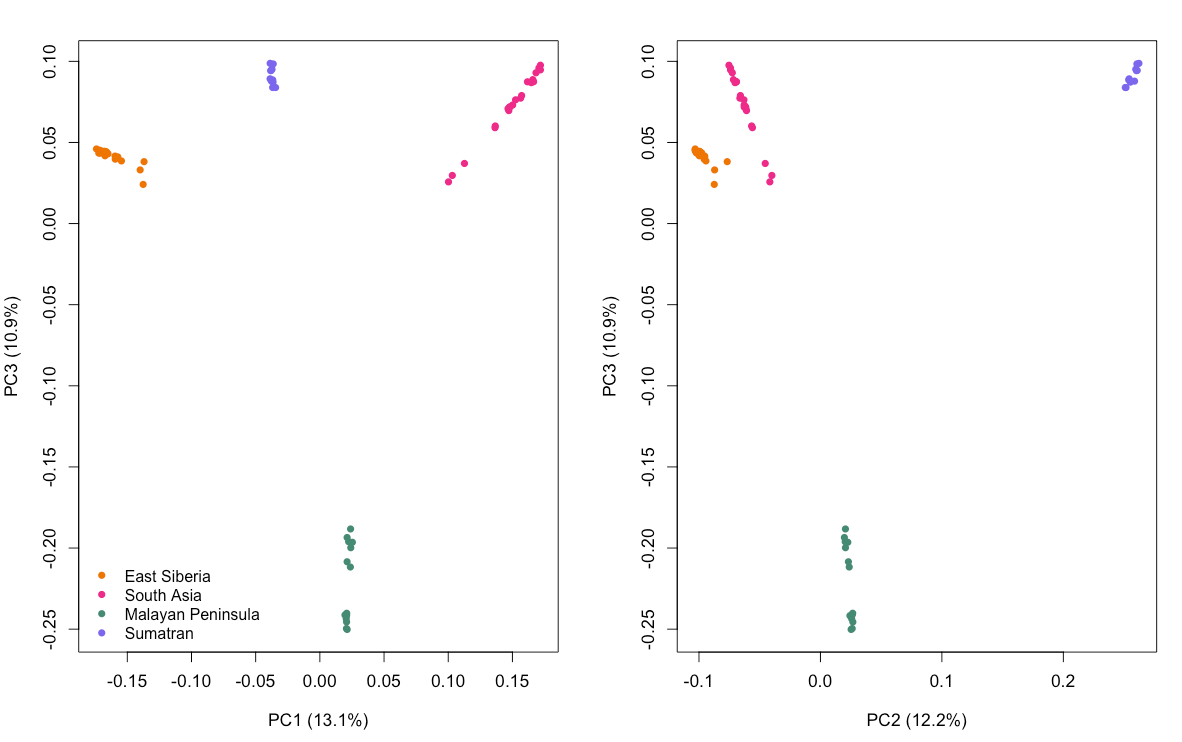
**Supplementary Figure 4**: Additional PCs (1-3) for all tigers as computed by PLINK2.

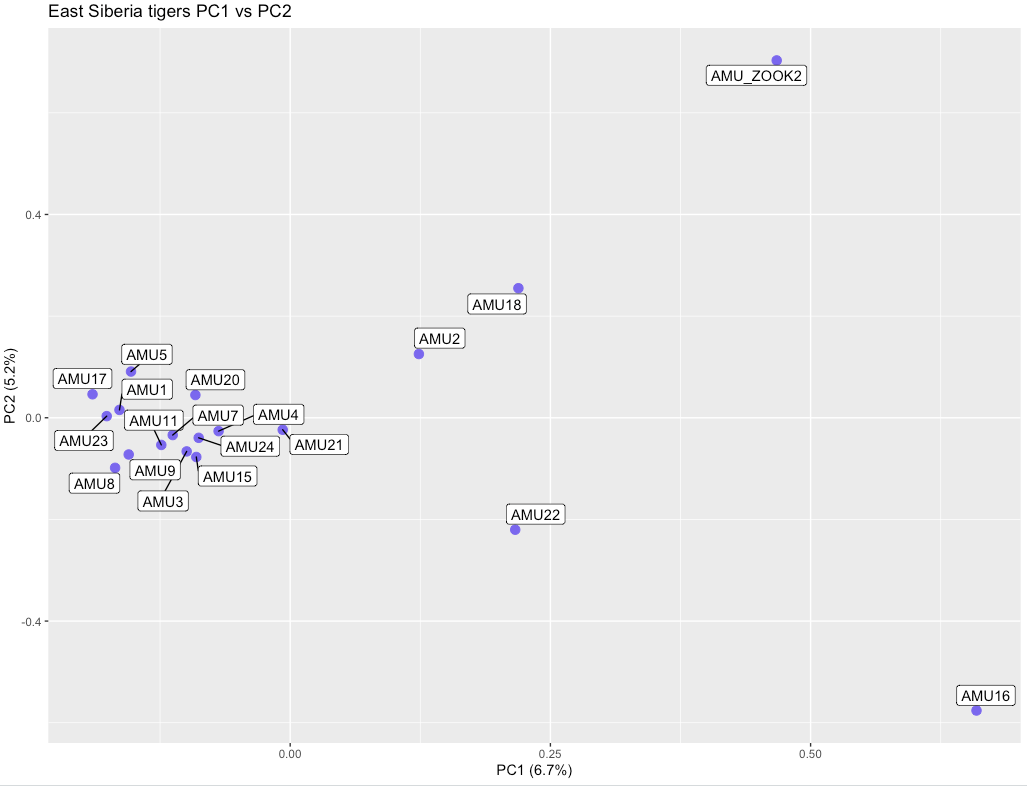

b.

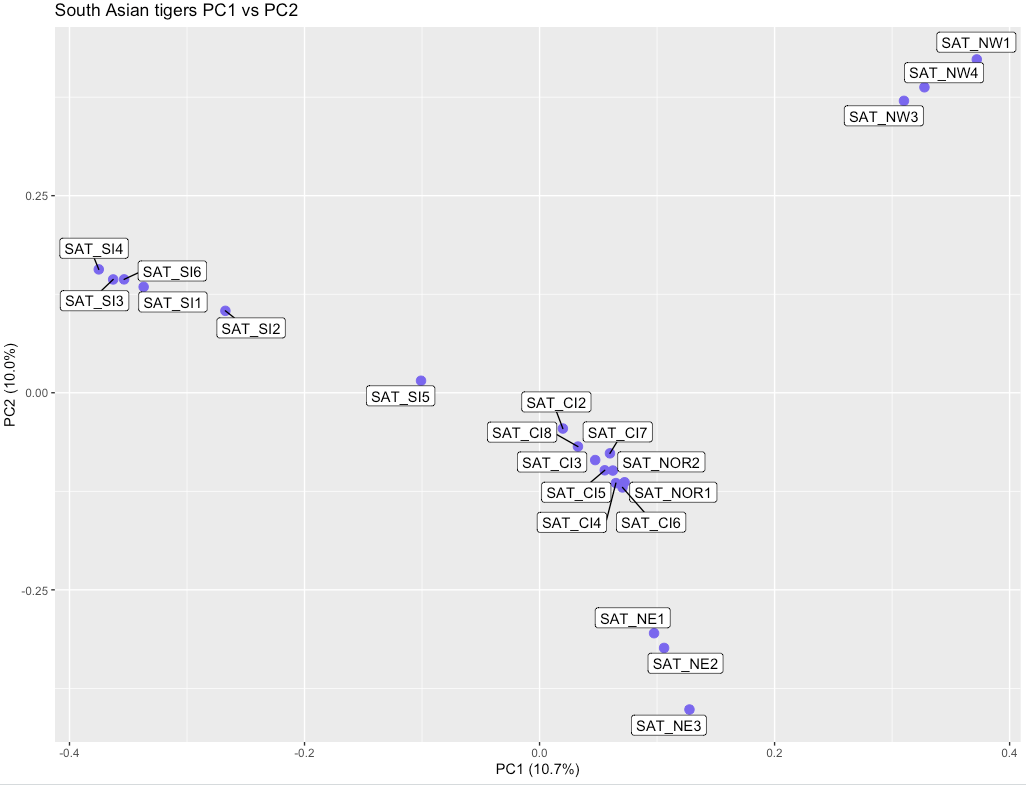

c.
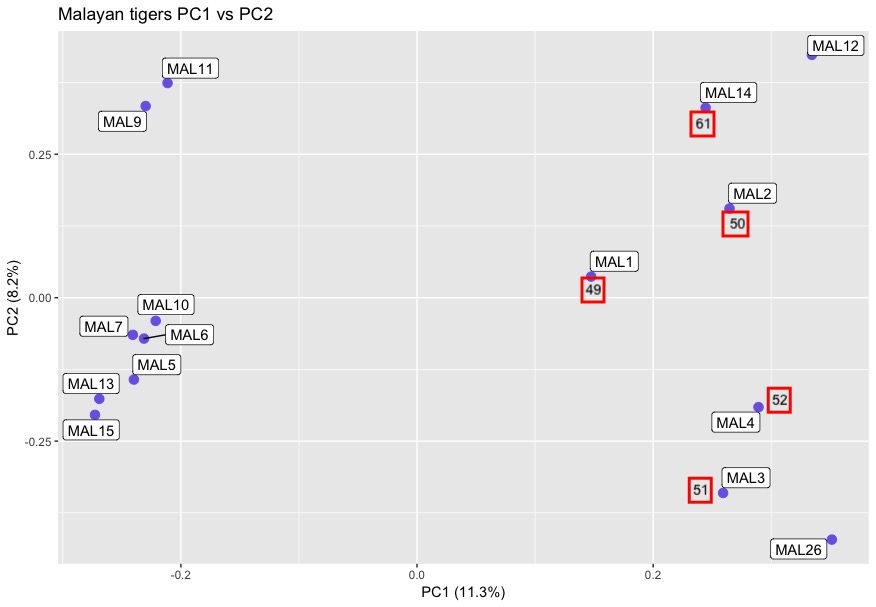

d.
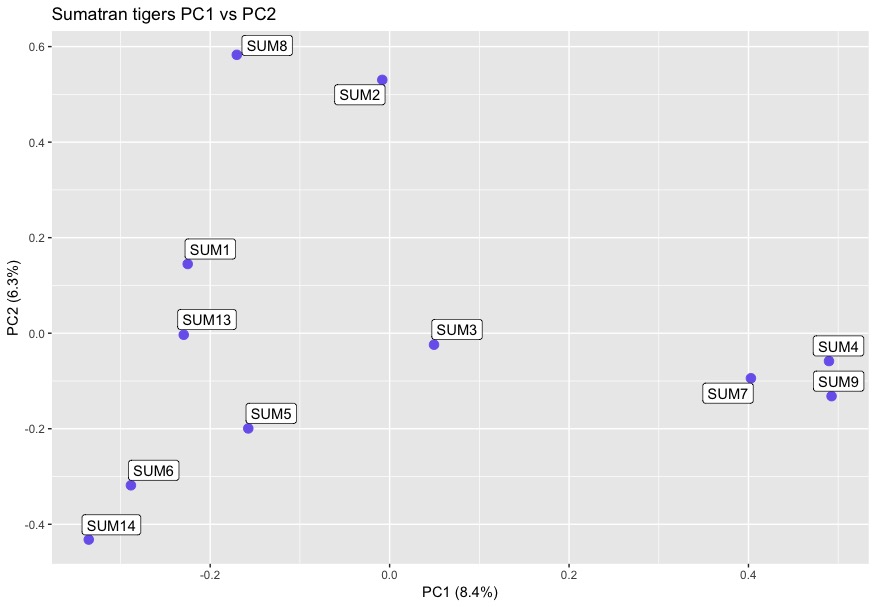

**Supplementary Figure 5 a, b, c, d**: Within subpopulation PCA for Amur, South Asian, Malayan, and Sumatran populations as computed by VCFtools with individuals labeled.

a.

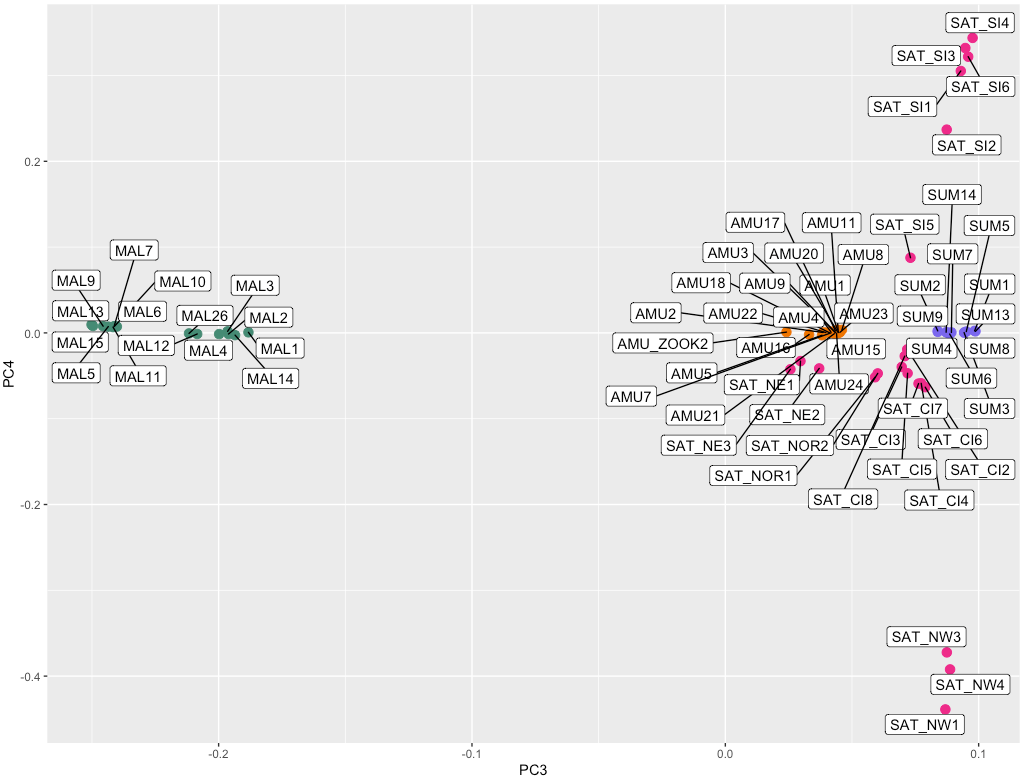

b.

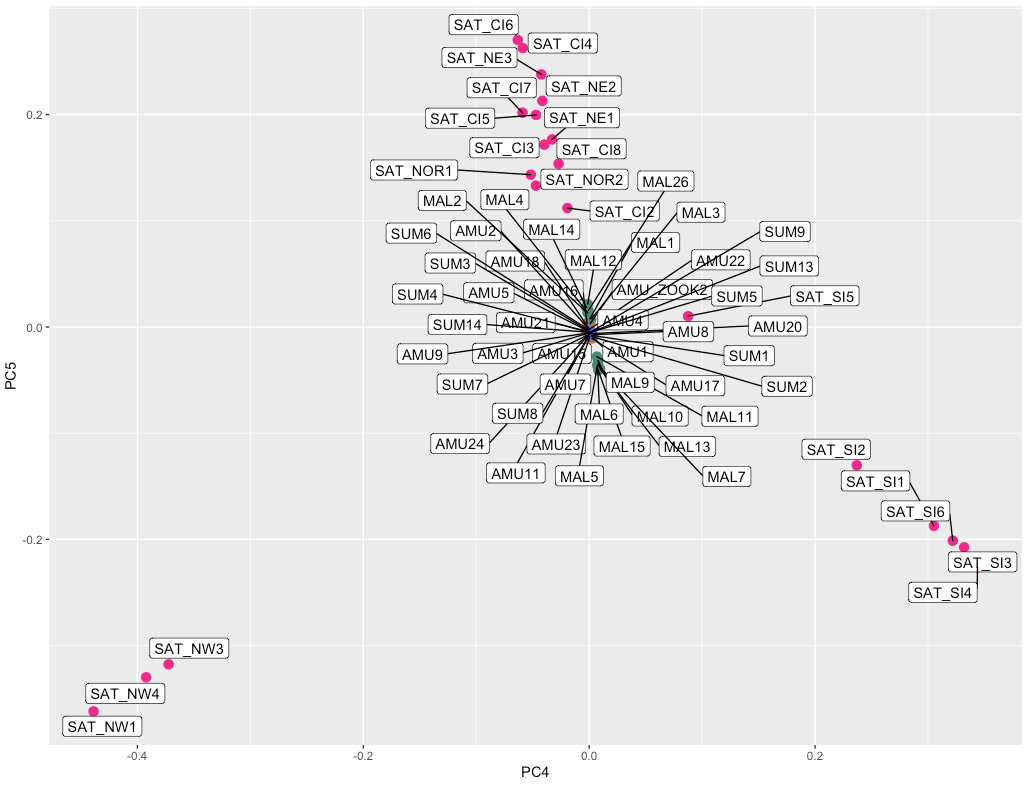

**Supplementary Figure 6**: PCA separation of South Asian and Malayan tiger populations across (a) PC axis 3 vs 4 and (b) PC axis 4 vs 5.

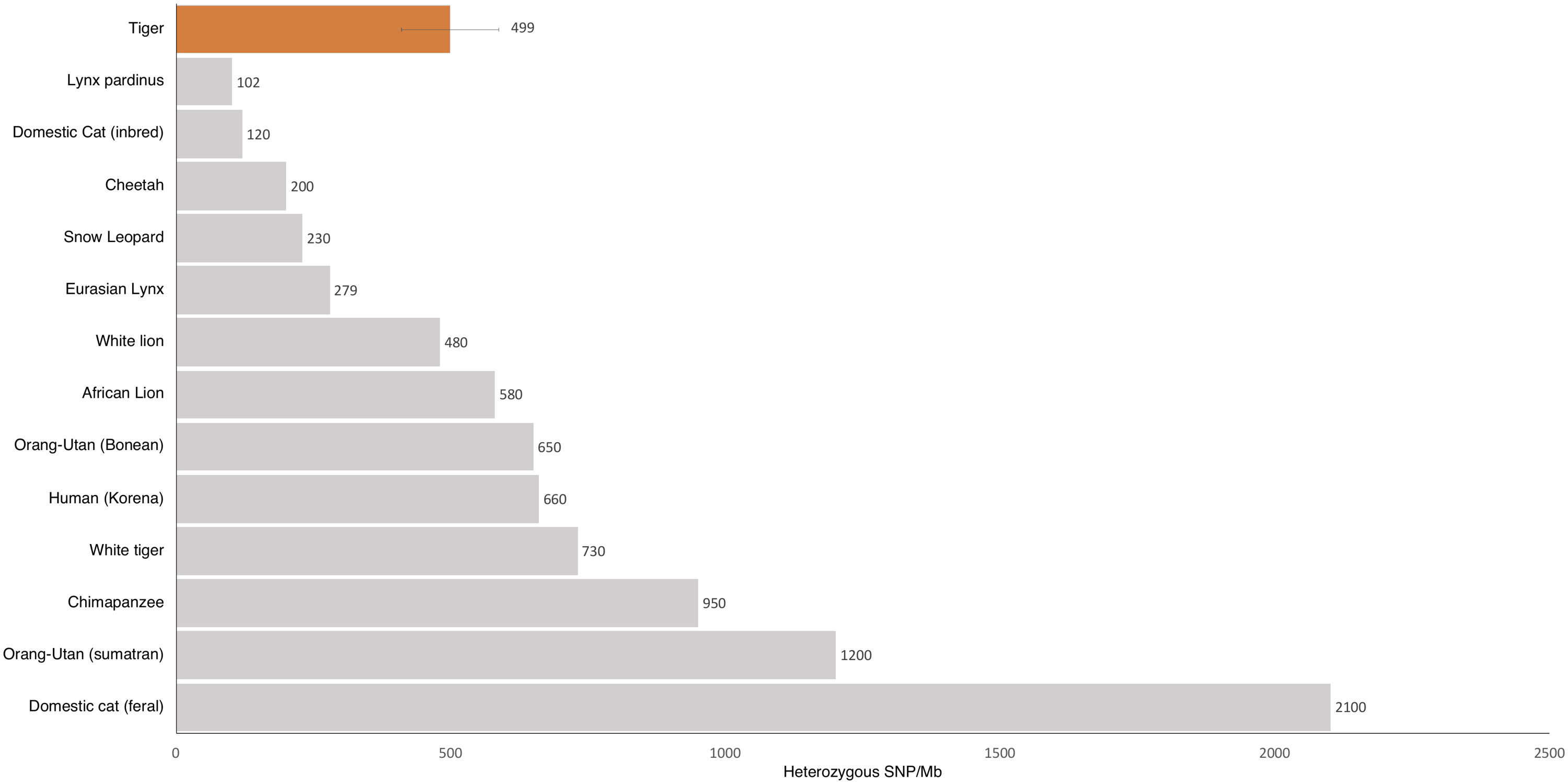

**Supplementary Figure 7**: Encounter rate for heterozygous SNVs as across the genome for various species. For most species, these are based on published estimates from single genomes. The error bar for the four tiger subspecies represented standard deviation across the mean.

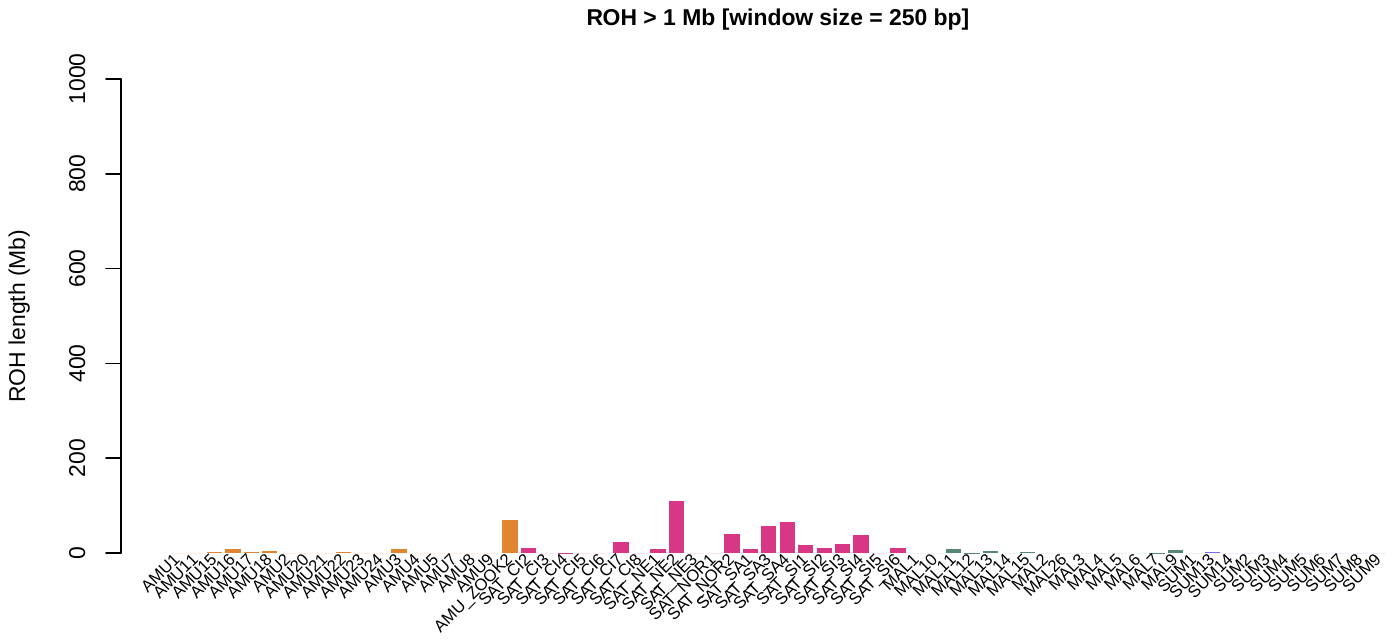

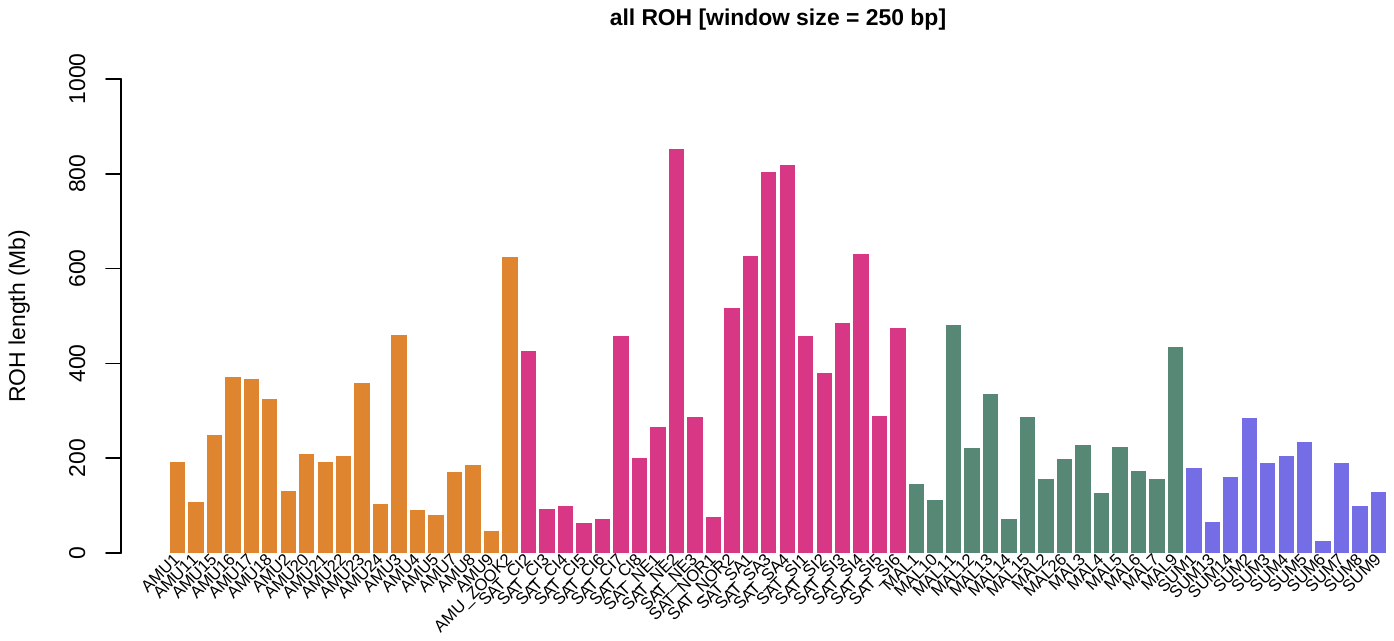

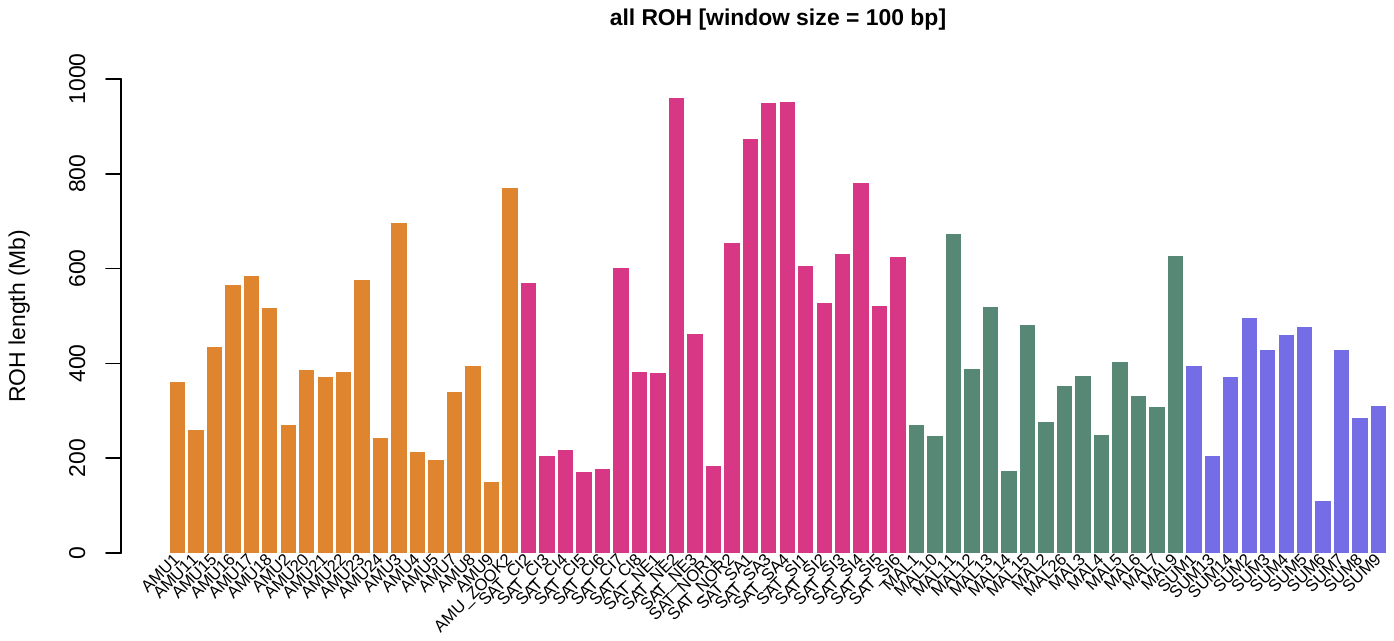

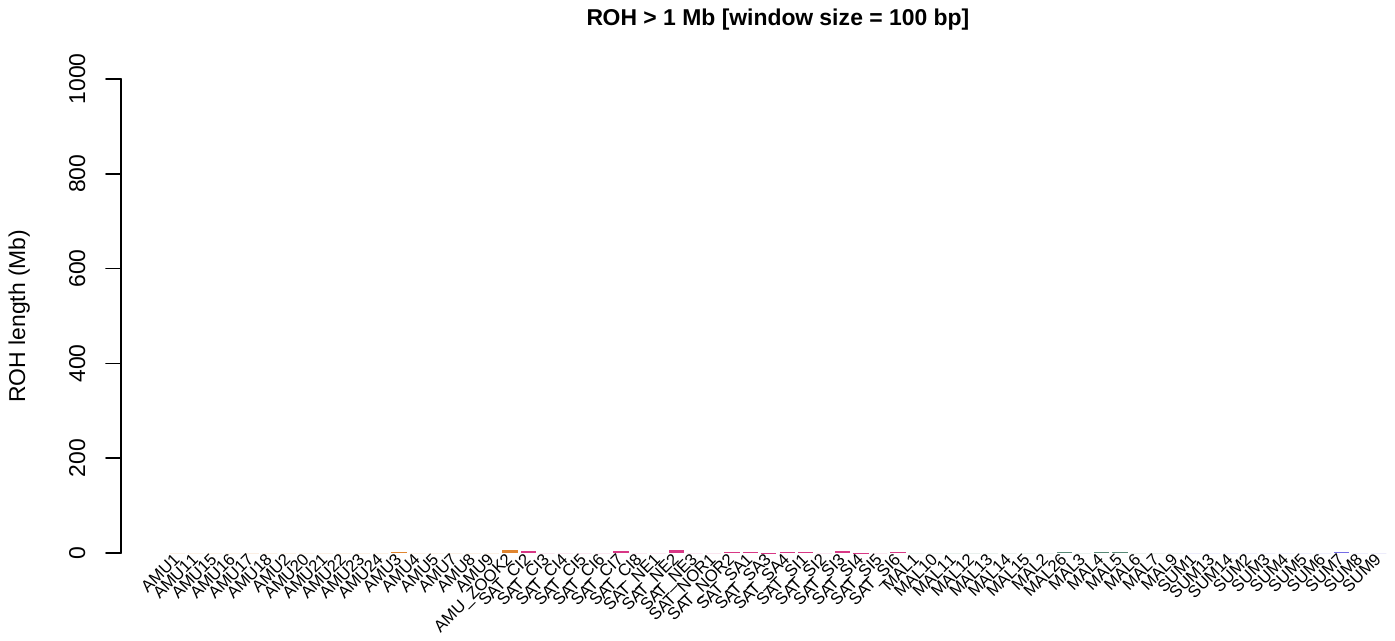

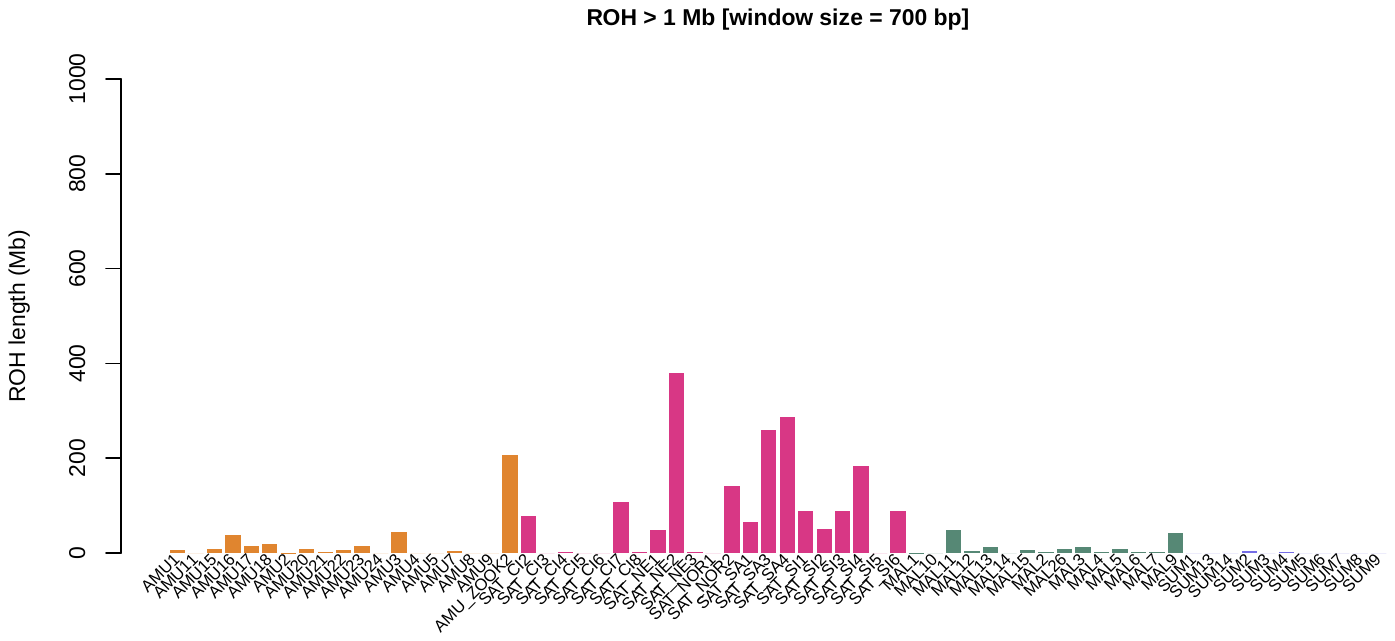

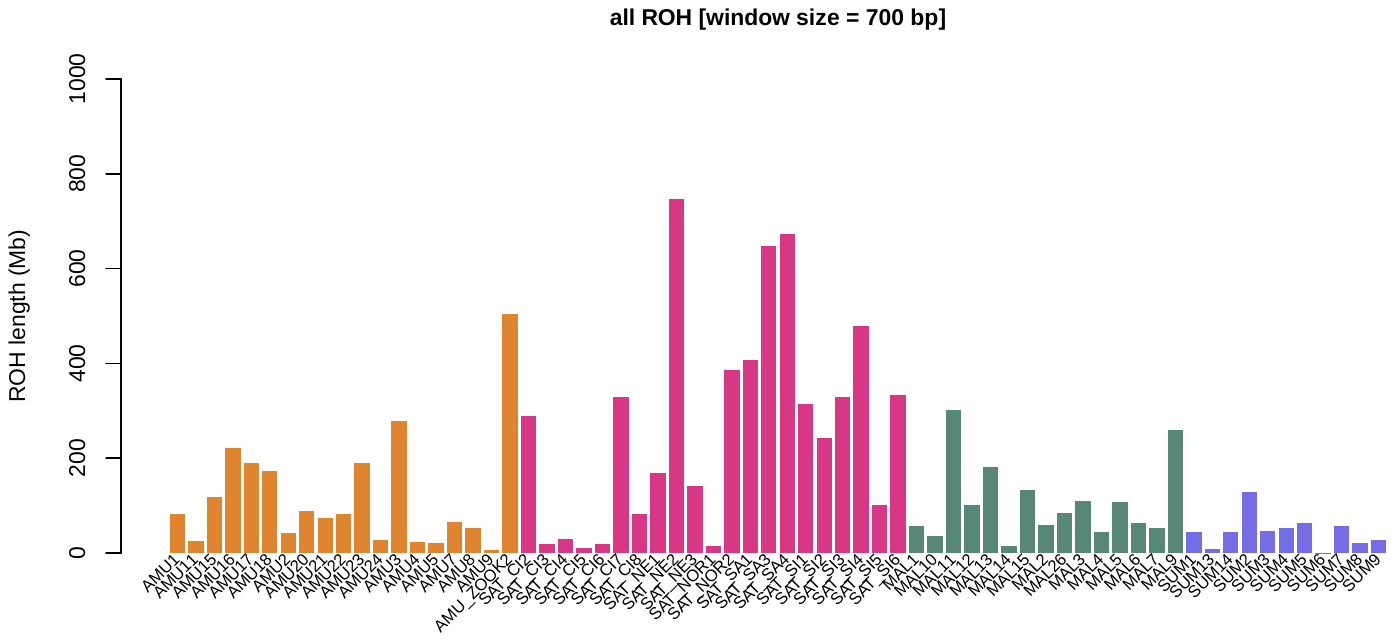

**Supplementary Figure 8**: Runs of Homozygosity predicted by GARLIC (Szpiech et al. 2017).

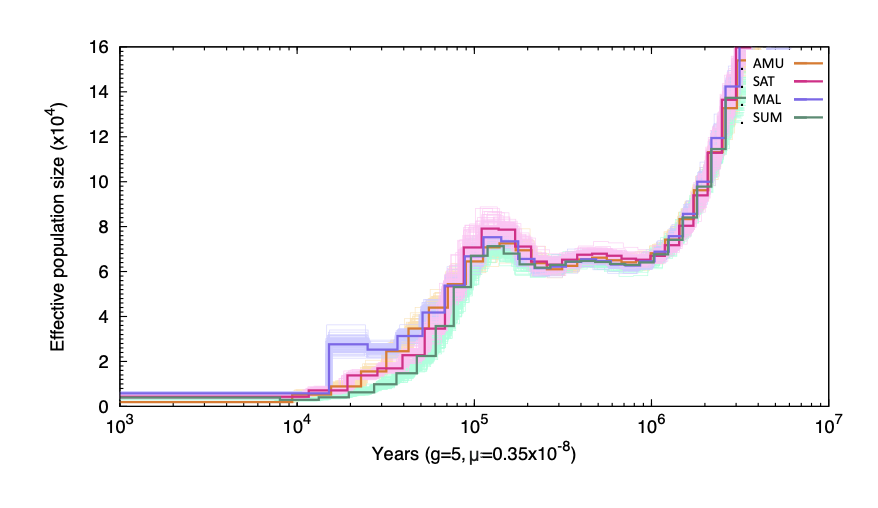

**Supplementary Figure 9**: PSMC plot for a single high coverage individual from each subspecies. Our plot is consistent with that observed by Liu et al (2018).

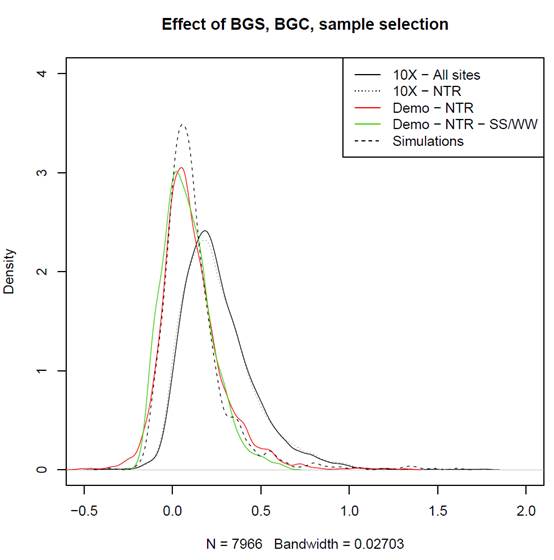

**Supplementary Figure 10**: Effect of background selection, biased gene conversion and sample selection on mPBS distributions in Siberian tigers. The black solid line represents mPBS values obtained when considering all sites and all individuals with average coverage above 10X (“10X – All sites”). The dashed thin black line corresponds to the same individuals but excluding transcribed regions and 50 kb flanking regions (“10X – NTR”). The red curve is computed using the same individuals that were used for demographic inference in non-transcribed regions (“Demo – NTR”). The green curve is obtained after filtering out sites potentially affected by biased gene conversion (“Demo – NTR – SS/WW”). Finally, the dashed black line corresponds to simulated values under the demographic model inferred from NTR-SS/WW sites (“Simulations”).

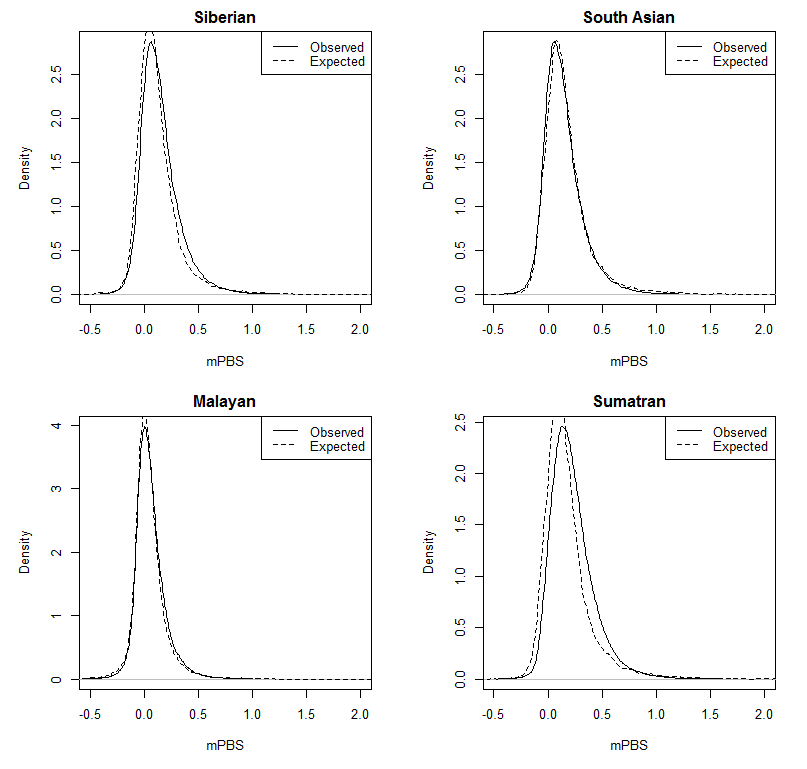

**Supplementary Figure 11**: Observed and expected mPBS distributions. For each population, we represented the observed against expected mPBS values obtained using simulations under a purely neutral model based on inferred demographic parameters. mPBS values have been computed using the same individuals as for demographic inference (Supplementary Table 7, Dataset 1).

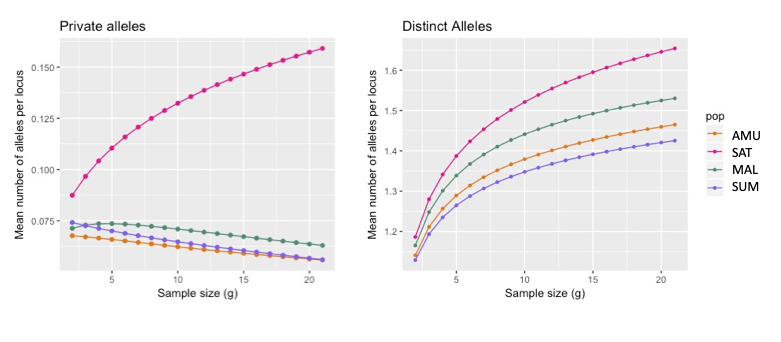

**Supplementary Figure 12**: ADZE plots of private alleles (alleles not found in other populations; left panel) and distinct (alleles unique to any given population, but allowing overlaps between populations).
**Supplementary tables**

**Supplementary Table 1**: List of tiger samples used within the coverage, in addition to sequencing information and associated AZA studbook numbers if associated.

| Map number | Sample ID | % bases covered | average depth | Source | Sample courtesy | SSB number | ancestry/location |
| --- | --- | --- | --- | --- | --- | --- | --- |
| 1 | AMU_ZOOK2 | 99 | 26.8784 | Captive | Published |  |  |
| 2 | AMU1 | 100 | 27.6484 | Captive | Omaha Zoological Society | 4203 | F1 |
| 3 | AMU2 | 100 | 26.6904 | Captive | San Diego Zoo | 260 | F1 |
| 4 | AMU3 | 91 | 4.43384 | Captive | Omaha Zoological Society | 3957 | F1 |
| 5 | AMU4 | 100 | 22.0467 | Wild | WCS Russia Program |  | Nezhinka river, Primorskii |
| 6 | AMU5 | 99 | 17.6047 | Wild | WCS Russia Program |  | Vyazemsky, Khabarovskii |
| 7 | AMU7 | 99 | 8.41758 | Wild | WCS Russia Program |  | Solontsovyi village, Khabarovskii |
| 8 | AMU8 | 99 | 14.387 | Wild | WCS Russia Program |  | Blagodatnoye, Primorskii |
| 9 | AMU9 | 100 | 19.2591 | Wild | WCS Russia Program |  | Blagodatnoye, Primorskii |
| 10 | AMU11 | 99 | 18.1544 | Wild | WCS Russia Program |  | Kunaleyka, Primorskii |
| 11 | AMU15 | 97 | 5.7804 | Captive | San Francisco Zoo | 4500 | F2 |
| 12 | AMU16 | 94 | 4.55855 | Captive | San Diego Zoo | 2686 | F2 |
| 13 | AMU17 | 96 | 5.39823 | Captive | San Diego Zoo | 5049 | F2 |
| 14 | AMU18 | 97 | 5.48226 | Captive | San Diego Zoo | 2879 | F2 |
| 15 | AMU20 | 100 | 27.6484 | Captive | Omaha Zoological Society | 2429 | F1 |
| 16 | AMU21 | 98 | 8.18089 | Captive | Omaha Zoological Society | 4013 | F2 |
| 17 | AMU22 | 96 | 5.92546 | captive | Omaha Zoological Society | 4187 | F2 |
| 18 | AMU23 | 94 | 4.8623 | captive | San Diego Zoo | 4638 | F2 |
| 19 | AMU24 | 100 | 17.1433 | Wild | WCS Russia Program |  | Nezhinka river, Primorskii |
| 20 | SAT_CI2 | 99 | 8.7746 | Wild | CCMB, Anuradha Reddy |  | Chandrapur |
| 21 | SAT_CI3 | 99 | 23.6797 | Wild | Wildlife Institute of India |  | Kanha Tiger Reserve |
| 22 | SAT_CI4 | 99 | 14.378 | Wild | Wildlife Institute of India |  | Kanha Tiger Reserve |
| 23 | SAT_CI5 | 99 | 14.6175 | Wild | Wildlife Institute of India |  | Kanha Tiger Reserve |
| 24 | SAT_CI6 | 99 | 18.9152 | Wild | Wildlife Institute of India |  | Kanha Tiger Reserve |
| 25 | SAT_CI7 | 99 | 9.33063 | Wild | Wildlife Institute of India |  | Kanha Tiger Reserve |
| 26 | SAT_CI8 | 90 | 5.23045 | Wild | CCMB, Anuradha Reddy |  | Pench Tiger Reserve |
| 27 | SAT_NE1 | 100 | 21.5036 | Wild | Aranyak |  | Assam |
| 28 | SAT_NE2 | 100 | 22.5455 | wild | Aranyak |  | Assam |
| 29 | SAT_NE3 | 99 | 14.8182 | wild | Aranyak |  | Assam |
| 30 | SAT_NOR1 | 99 | 22.7056 | wild | Wildlife Institute of India |  | Corbett Tiger Reserve |
| 31 | SAT_NOR2 | 100 | 23.6836 | wild | Wildlife Institute of India |  | Corbett Tiger Reserve |
| 32 | SAT_NW1 | 83 | 5.56259 | wild | Wildlife Institute of India |  | Ranthambore Tiger Reserve |
| 33 | SAT_NW3 | 100 | 21.6486 | wild | NCBS |  | Ranthambore Tiger Reserve |
| 34 | SAT_NW4 | 99 | 24.0216 | wild | NCBS |  | Ranthambore Tiger Reserve |
| 35 | SAT_SI1 | 100 | 20.5914 | wild | NCBS |  | Bandipur Tiger Reserve |
| 36 | SAT_SI2 | 99 | 9.14737 | wild | NCBS |  | Waynad Wildlife Sanctuary |
| 37 | SAT_SI3 | 100 | 21.6486 | wild | Kerala Forest Department |  | Waynad Wildlife Sanctuary |
| 38 | SAT_SI4 | 99 | 24.0216 | wild | Kerala Forest Department |  | Waynad Wildlife Sanctuary |
| 39 | SAT_SI5 | 99 | 13.0038 | wild | Kerala Forest Department |  | Periyar Tiger Reserve |
| 40 | SAT_SI6 | 100 | 18.3987 | wild | Kerala Forest Department |  | Waynad Wildlife Sanctuary |
| 41 | MAL1 | 100 | 32.8678 | wild caught | Omaha Zoological Society | 149 | Wild caught, Terenggan, Paka |
| 42 | MAL2 | 100 | 31.7846 | wild caught | El Paso zoo | 194 | Wild caught, Kelantan |
| 43 | MAL3 | 100 | 29.5029 | wild caught | San Diego Zoo | 196 | Wild caught, Kelantan |
| 44 | MAL4 | 100 | 21.9693 | wild caught | San Diego Zoo | 220 | Wild caught, Jeli |
| 45 | MAL5 | 93 | 4.19328 |  | San Diego Zoo | 37 | F1 |
| 46 | MAL6 | 96 | 4.74293 |  | San Diego Zoo | 184 | F2 |
| 47 | MAL7 | 95 | 4.99761 |  | WCS, Bronx Zoo | 190 | F2, sibling of 184 |
| 48 | MAL9 | 96 | 5.39176 |  | Omaha Zoological Society | 173 | F1, sibling of 66 |
| 49 | MAL10 | 97 | 6.18214 |  | San Diego Zoo | 168 | F2, sibling of 184 |
| 50 | MAL11 | 95 | 5.01651 |  | San Diego Zoo | 212 | F2 |
| 51 | MAL12 | 94 | 4.92575 |  | Omaha Zoological Society | 214 | F2, sibling of 212 |
| 52 | MAL13 | 96 | 5.91426 |  | San Diego Zoo | 66 | F1, sibling of 37 |
| 53 | MAL14 | 99 | 16.2318 |  | Omaha Zoological Society | 137 | Wild caught, Pehang |
| 54 | MAL15 | 97 | 6.14838 |  | San Diego Zoo | 67 | F2, sibling of 212 |
| 55 | MAL26 | 95 | 5.30186 |  | San Diego Zoo | 244 | F1 |
| 56 | SUM1 | 100 | 30.5978 |  | Omaha Zoological Society | 527 | F1 |
| 57 | SUM2 | 100 | 31.2564 |  | Omaha Zoological Society | 564 | F2 |
| 58 | SUM3 | 96 | 5.24071 |  | San Francisco Zoo | 722 | F2 |
| 59 | SUM4 | 91 | 4.45531 |  | San Diego Zoo | 311 | F2 |
| 60 | SUM5 | 95 | 4.74431 |  | San Diego Zoo | 381 | F2 |
| 61 | SUM6 | 100 | 21.6518 |  | San Diego Zoo | 1282 | F2 |
| 62 | SUM7 | 95 | 4.92926 |  | San Diego Zoo | 316 | F2, sibling of 311 |
| 63 | SUM8 | 97 | 6.3304 |  | San Diego Zoo | 1110 | F2 |
| 64 | SUM9 | 97 | 6.38643 |  | San Diego Zoo | 312 | F2, sibling of 311 |
| 65 | SUM13 | 99 | 14.8678 |  | San Diego Zoo | 768 | F2 |
| 66 | SUM14 | 96 | 5.75034 |  | San Diego Zoo | 1281 | F2, sibling of 1282 |

**Supplementary Table 2**: Assemblathon-2 statistics of new genome compared with PanTig1.0 (Cho et al., 2013).

|  | Belahat 10x (Maltig1.0) | PanTig1.0 |
| --- | --- | --- |
| Contig N50 | 1.8 Mb | 39 kb |
| Contig L90 | 13,816 | 62,406 |
| Scaffold N50 | 21.3 Mb | 8.9 Mb |
| Scaffold L90 | 109 | 284 |
| % Scaffold Ns | 1.31% | 2.44% |
| Number of contigs | 36,444 | 126,370 |
| Number of scaffolds | 10,077 | 1,479 |

**Supplementary Table 3**: Details of analyses and samples used

| Analysis | SNP calling method | SNP Filters | Individuals used |
| --- | --- | --- | --- |
| **Population Clustering** |  |  |  |
| ADMIXTURE | freebayes | Q=30, GQ=30, DP=10, maf=0.025, max_miss=5% | ALL |
| PCA | freebayes | Q=30, GQ=30, DP=10, maf=0.025, max_miss=5% | ALL |
| NetStruct | freebayes | Q=30, GQ=30, DP=10, maf=0.025, max_miss=5% | ALL |
| FST | freebayes | Q=30, GQ=30, DP=10, maf=0.025, max_miss=5% | ALL |
| **Genetic Diversity** |  |  |  |
| SNV encounter rate | freebayes | Q=30, GQ=30, DP=10, maf=0.025, max_miss=5% | SAT_SI6, SAT_SI4, SAT_SI3, SAT_SI1, SAT_NW4, SAT_NW3, SAT_NOR1, SAT_NOR2, SAT_NE1, SAT_NE2, SAT_CI8, SAT_CI6, SAT_CI4, SAT_CI3, SUM13, SUM6, SUM2, SUM1, MAL14, MAL4, MAL3, MAL2, MAL1, AMU24, AMU11, AMU9, AMU5, AMU4, AMU2, AMU1 |
| ADZE | freebayes | Q=30, GQ=30, DP=10, maf=0.025, max_miss=5% | ALL |
| Nucleotide diversity | freebayes | Q=30, GQ=30, DP=10, maf=0.025, max_miss=5% | ALL |
| **Demographic history** |  |  |  |
| PSMC | bcftools | DP=10, maxDP=2*(average depth) | AMU1, MAL1, SUM2, SAT_SI3 |
| FastSimcoal (Global demographic history) | freebayes | Q=30, GQ=30, DP=10, regions within 50Kb of transcribing regions removed | SUM2, SUM1, SUM6, SAT_SI3, SAT_SI4, SAT_SI1, MAL1, MAL2, MAL3, MAL4, AMU2, AMU1, AMU4 |
| FastSimcoal (South Asian tiger demographic history) | freebayes | Q=30, GQ=30, DP=10, regions within 50Kb of transcribing regions removed | MAL1, MAL2, MAL3, SAT_CI3, SAT_CI6, SAT_NE2, SAT_NE1, SAT_NW4, SAT_NW3, SAT_SI3, SAT_SI4, SAT_SI1 |
| **Selection scans** |  |  |  |
| mPBS | freebayes | Q=30, GQ=30, DP=10 | SAT_SI6, SAT_SI5, SAT_SI4, SAT_SI3, SAT_SI1, SAT_NW4, SAT_NW3, SAT_NOR1, SAT_NOR2, SAT_NE1, SAT_NE2, SAT_NE3, SAT_CI6, SAT_CI5, SAT_CI4, SAT_CI3, SUM13, SUM6, SUM2, SUM1, MAL14, MAL4, MAL3, MAL2, MAL1, AMU24, AMU20,AMU11,AMU9, AMU8, AMU5, AMU4, AMU2, AMU1, AMU_ZOOK2 |
| **Inbreeding** |  |  |  |
| bcftools ROH | freebayes | Q=30, GQ=30, DP=10, maf=0.025, max_miss=5%, noSex | SAT_SI6, SAT_SI5, SAT_SI4, SAT_SI3, SAT_SI1, SAT_NW4, SAT_NW3, SAT_NOR1, SAT_NOR2, SAT_NE1, SAT_NE2, SAT_NE3, SAT_CI8, SAT_CI6, SAT_CI5, SAT_CI4, SAT_CI3, SUM13, SUM6, SUM2, SUM1, MAL14, MAL4, MAL3, MAL2, MAL1, AMU24, AMU20, AMU9, AMU5, AMU4, AMU2, AMU1, AMU_ZOOK2 |
| GARLIC | freebayes | Q=30, GQ=30, DP=10, maf=0.025, max_miss=5%, noSex | ALL |

**Supplementary Table 4**: Weighted F_ST_ between subspecies as computed by VCFtools.

| Population | South Asian | Malayan | Sumatran |
| --- | --- | --- | --- |
| Amur | 0.200 | 0.230 | 0.318 |
| South Asian |  | 0.164 | 0.242 |
| Malayan |  |  | 0.280 |

**Supplementary Table 5**: Weighted F_ST_ between putative South Asian sub-populations as computed by VCFtools. NOR=North India, SI=South India, NW=Northwest India, CI=Central India, NE=Northeast India.

|  | SAT_NOR | SAT_SI | SAT_NW | SAT_CI |
| --- | --- | --- | --- | --- |
| SAT_NE | 0.12752 | 0.18127 | 0.27993 | 0.11281 |
| SAT_NOR |  | 0.17404 | 0.29921 | 0.09356 |
| SAT_SI |  |  | 0.27524 | 0.12259 |
| SAT_NW |  |  |  | 0.20262 |

**Supplementary Table 6**: Selected samples and number of retained sites for the two datasets used in demographic inference.

|  | # SUM  inds | # SAT  inds | # MAL  inds | # AMU  inds | Samples | # polymorphic sites without filtering | # polymorphic sites after filtering |
| --- | --- | --- | --- | --- | --- | --- | --- |
| Dataset 1 | 3 | 3 | 4 | 3 | SUM2, SUM1, SUM6,  SAT_SI3, SAT_SI4, SAT_SI1, MAL1, MAL2, MAL3, MAL4, AMU2, AMU1, AMU4 | 2,892,663 | 265,560 |
| Dataset 2 | 0 | 9 | 3 | 0 | MAL1, MAL2, MAL3, SAT_CI3, SAT_CI6, SAT_NE2, SAT_NE1, SAT_SA4, SAT_SA3, SAT_SI3, SAT_SI4, SAT_SI1 | 2,935,205 | 267,933 |

**Supplementary Table 7**: Estimated demographic parameters and confidence intervals for the modeling of the four subspecies differentiation from an Asian metapopulation

| **Parameter acronyms** | **Parameter descriptions** | **Maximum likelihood estimate** | **lower  limit CI95%** | **upper limit CI95%** |
| --- | --- | --- | --- | --- |
| N_ANCEST | Ancestral Asian metapopulation size | 112,088 | 106,999 | 128,083 |
| N_ASIA | Asian metapopulation size | 8,281 | 6,443 | 15,953 |
| N_SUM | Sumatra population size | 5,380 | 3,765 | 11,892 |
| N_SAT | South Asian population size | 5,606 | 4,395 | 11,866 |
| N_MAL | Malayan population size | 12,801 | 7,338 | 13,392 |
| N_AMU | Amur population size | 9,735 | 3692 | 12872 |
| TD_SUM | Sumatran divergence time | 9,147 | 6,378 | 9,650 |
| TD_SAT | South Asian divergence time | 11,842 | 9,900 | 13,795 |
| TD_MAL | Malayan divergence time | 7,775 | 2,820 | 8,490 |
| TD_AMU | Amur divergence time | 7,562 | 3,248 | 8,118 |
| IB_SUM | Sumatran initial bottleneck (founder effect) | 0.1241 | 0.0427 | 0.2329 |
| IB_SAT | South Asian initial bottleneck (founder effect) | 0.0827 | 0.0024 | 0.0987 |
| IB_MAL | Malayan initial bottleneck (founder effect) | 0.0032 | 0.0015 | 0.0179 |
| IB_AMU | Amur initial bottleneck (founder effect) | 0.0153 | 0.0023 | 0.0514 |
| RB_SUM | Sumatran recent bottleneck | 0.2035 | 0.1044 | 0.2959 |
| RB_SAT | South Asian recent bottleneck | 0.3106 | 0.3006 | 0.3777 |
| RB_MAL | Malayan recent bottleneck | 0.1204 | 0.0844 | 0.1457 |
| RB_AMU | Amur recent bottleneck | 0.2372 | 0.1768 | 0.2726 |
| Nm_SUM | SUM immigration rate | 0.1137 | 0.0004 | 0.2862 |
| Nm_SAT | SAT immigration rate | 3.1138 | 2.5340 | 4.7430 |
| Nm_MAL | MAL immigration rate | 0.0118 | 0.0008 | 1.0269 |
| Nm_AMU | AMU immigration rate | 0.2040 | 0.0008 | 1.1302 |
| IB_ANC | Ancestral Asian bottleneck | 1.0684 | 1.0156 | 1.2742 |
| T_BOT | Time of ancestral bottleneck | 234,238 | 196,525 | 245,795 |

Population sizes (N) are in number of diploid individuals assuming a mutation rate of 0.35e-8 (Liu et al. 2018). Times (T) are in years assuming a generation time of 5 years. Immigration rates (Nm) values are expressed as twice the number of diploids entering the population per generation, coming from the Asian metapopulation. Bottlenecks intensities (IB and RB) are indicated as bottleneck duration in units of 2N generations, but they were modeled as instantaneous bottlenecks (see fastsimcoal manual http://cmpg.unibe.ch/software/fastsimcoal2/man/fastsimcoal26.pdf, p. 27).

**Supplementary Table 8**: Estimated demographic parameters and confidence intervals for the modeling of four South Asian population differentiation from an Indian metapopulation

| **Parameter acronyms** | **Parameter descriptions** | **Maximum likelihood estimate** | **lower limit CI95%** | **upper limit CI95%** |
| --- | --- | --- | --- | --- |
| N_ANCEST | Ancestral Asian metapopulation size | 122,089 | 120,570 | 123,548 |
| N_META_IND | Indian metapopulation size | 7,987 | 7,459 | 11,208 |
| N_SAT_CI | CI population size | 4,481 | 3,099 | 11,440 |
| N_SAT_NE | NE population size | 11,525 | 5,208 | 13,160 |
| N_SAT_NW | NW population size | 4,742 | 2,026 | 11,575 |
| TD_SAT_CI | CI divergence time | 2,040 | 999 | 2,986 |
| TD_SAT_NE | NE divergence time | 6,784 | 5,924 | 8,109 |
| TD_SAT_NW | NW divergence time | 1,186 | 935 | 2,541 |
| TD_SAT_SI | SI divergence time | 1,548 | 971 | 2,429 |
| TD_SAT | Indian metapopulation divergence time | 8,462 | 8,056 | 9,110 |
| IB_SAT_CI | CI initial bottleneck (founder effect) | 0.0056 | 0.0025 | 0.0381 |
| IB_SAT_NE | NE initial bottleneck | 0.0166 | 0.0018 | 0.0644 |
| IB_SAT_SA | NW initial bottleneck | 0.3110 | 0.1137 | 0.4939 |
| IB_SAT_SI | SI initial bottleneck | 0.0078 | 0.0014 | 0.0119 |
| IB_META_IND | Indian metapopulation initial bottleneck | 0.0527 | 0.0364 | 0.0674 |
| RB_SAT_CI | CI recent bottleneck | 0.0042 | 0.0021 | 0.0375 |
| RB_SAT_NE | NE recent bottleneck | 0.0598 | 0.0028 | 0.0836 |
| RB_SAT_NW | NW recent bottleneck | 0.2076 | 0.0189 | 0.4141 |
| NM__Asia2IND | Indian immigration rate (from Asia) | 0.2031 | 0.0004 | 0.3015 |
| NM__SAT_CI | CI immigration rate | 0.0049 | 0.0010 | 1.1668 |
| NM__SAT_NE | NE immigration rate | 0.0104 | 0.0004 | 0.6793 |
| NM__SAT_NW | NW immigration rate | 0.2746 | 0.0004 | 0.7654 |
| NM__SAT_SI | SI immigration rate | 0.0103 | 0.0005 | 1.4335 |

Population sizes (N) are in number of diploid individuals, assuming a mutation rate of 0.35e-8 (Liu et al. 2018). Times (T) are in years, assuming a generation time of 5 years. Immigration rates (Nm) values are expressed as twice the number of diploids entering the population per generation, coming from the Asian metapopulation. Bottlenecks intensities (IB and RB) are indicated as bottleneck duration in units of 2N generations, but they were modeled as instantaneous bottlenecks (see fastsimcoal manual http://cmpg.unibe.ch/software/fastsimcoal2/man/fastsimcoal26.pdf, p. 27). The values of the following parameters SAT_SI were fixed to previously estimated values: N_SAT_SI=5606, RB_SAT_SI=0.3106.

**Supplementary Table 9**: Gene Ontology enrichment results. The 20 most significant GO terms are presented, as well as their total number of genes, the number of observed significant genes in a given term, the expected number of significant genes, the fold enrichment and the p-value of the Fisher’s exact test.

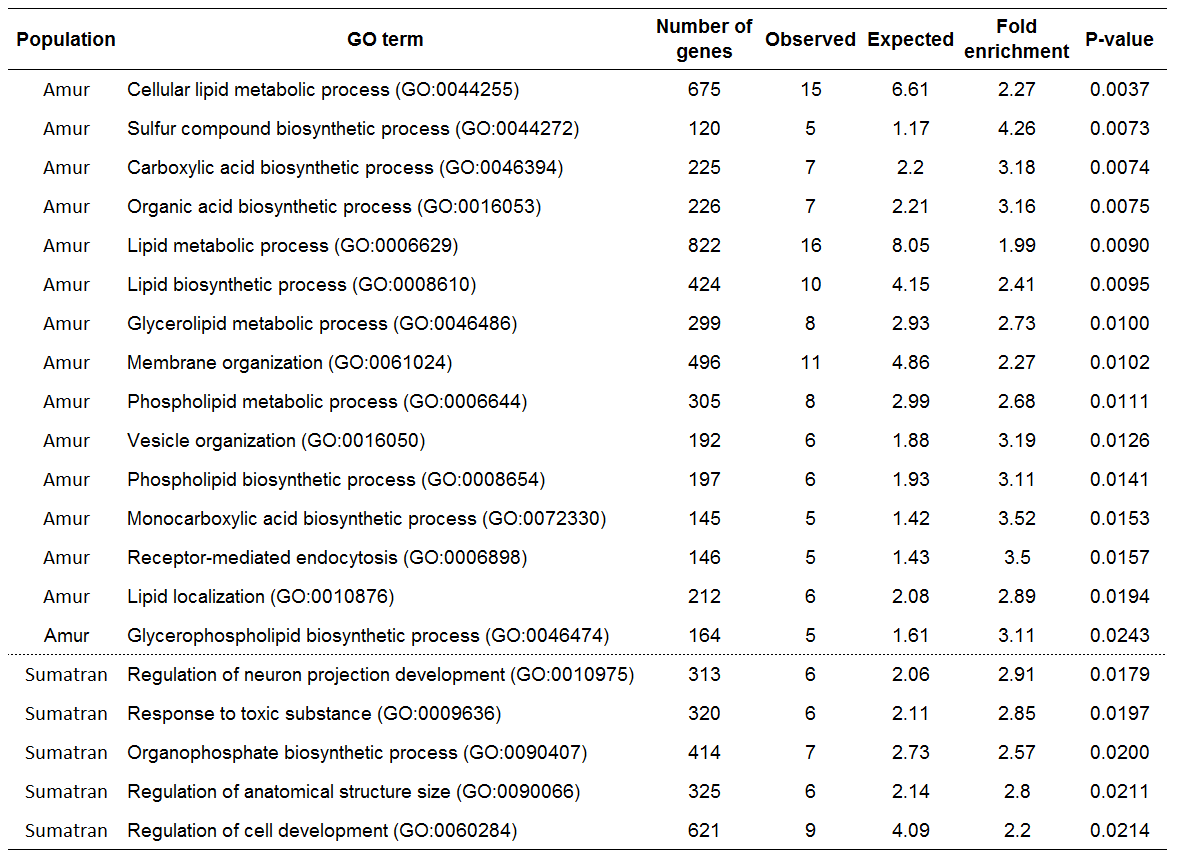

**Supplemental files**

Fastsimcoal input files used to estimate the Asian tiger population demography

TIGERS8e.tpl

```
//Simple model of tiger differentiation, no bottleneck
5  samples to simulate
//Population effective sizes
N_SUM
N_SAT
N_MAL
N_AMU
N_GHOST
//Samples sizes
6
6
8
6
0
//Growth rates : negative growth implies population expansion
0
0
0
0
0
//Number of migration matrices : 0 implies no migration between demes
2
//Migration matrix 0
0 0 0 0 0
0 0 0 0 0
0 0 0 0 0
0 0 0 0 0
0 0 0 0 0
//Migration matrix 1
0 0 0 0 MIG_SUM
0 0 0 0 MIG_SAT
0 0 0 0 MIG_MAL
0 0 0 0 MIG_AMU
0 0 0 0 0
//historical event: time, source, sink, migrants, new deme size, new growth rate, migration matrix index
16
0 0 0 0 1 0 0
50 0 0 0 RB_SUM 0 0 instbot
50 1 1 0 RB_SAT 0 0 instbot
50 2 2 0 RB_MAL 0 0 instbot
50 3 3 0 RB_AMU 0 0 instbot
50 0 0 0 1 0 1
TD_MAL 2 2 0 IB_MAL     0 1 instbot
TD_MAL 2 4 1 1          0 1
TD_AMU 3 3 0 IB_AMU     0 1 instbot
TD_AMU 3 4 1 1          0 1
TD_SAT 1 1 0 IB_SAT     0 1 instbot
TD_SAT 1 4 1 1          0 1
TD_SUM 0 0 0 IB_SUM     0 1 instbot
TD_SUM 0 4 1 1          0 1
TDMAX  4 4 0 R_ANC      0 0              //Ancestral population size change after last divergence
TANCBOT 4 4 0 IB_ANC    0 0 instbot      //Ancestral inst bot possible speciation...
//Number of independent loci [chromosome]
1 0
//Per chromosome: Number of contiguous linkage Block: a block is a set of contiguous loci
1
//per Block:data type, number of loci, per generation recombination and mutation rates and optional parameters
FREQ 1 0 1e-8 OUTEXP
```

TIGERS9B3Mv1_r8e.est

```
// Priors and rules file
// *********************

[PARAMETERS]
//#isInt? #name   #dist.#min  #max
//all Ns are in number of haploid individuals

//Population sizes -----------------------------------------
1  N_ANCEST      unif  10000 100000  output REFERENCE //Ancestral size (meta pop size as well)
1  N_GHOST       unif  1000  100000 output
1  N_SUM         unif  1000   10000  output bounded
1  N_SAT         unif  1000   10000  output bounded
1  N_MAL         unif  1000   10000  output bounded
1  N_AMU         unif  1000   10000  output bounded

//Divergence times -----------------------------------------
1  TD_SUM        unif  400 1000  output bounded
1  TD_SAT        unif  100 1000  output bounded
1  TD_MAL        unif  100 1000  output bounded
1  TD_AMU        unif  100 1000  output bounded

//Bottleneck intensities -----------------------------------
0  IB_SUM        logunif  0.001 1  output
0  IB_SAT        logunif  0.001 1  output
0  IB_MAL        logunif  0.001 1  output
0  IB_AMU        logunif  0.001 1  output

0  RB_SUM        unif  0.01 1  output
0  RB_SAT        unif  0.01 1  output
0  RB_MAL        unif  0.01 1  output
0  RB_AMU        unif  0.01 1  output

0  IB_ANC        logunif  0.001 2  output

//Gene flow rates ------------------------------------------
0 Nm_SUM     logunif 1e-4 5 output bounded
0 Nm_SAT     logunif 1e-4 5 output bounded
0 Nm_MAL     logunif 1e-4 5 output bounded
0 Nm_AMU     logunif 1e-4 5 output bounded

//Ancestral population size change ------------------------
1  TPLUSANCBOT    unif  1e3  1e5  hide

[RULES]

[COMPLEX PARAMETERS]

//Ancestral population size change ------------------------
1 MAX1 = TD_SUM%max%TD_SAT     hide
1 MAX2 = MAX1%max%TD_MAL       hide
1 TDMAX = MAX2%max%TD_AMU      hide
1 TANCBOT = TDMAX+TPLUSANCBOT  output

//Ancestral resize
0 R_ANC = N_ANCEST/N_GHOST     hide
0 R_PREV = N_PREV/N_ANCEST     hide

//Migration rates ------------------------------------------
0 MIG_SUM = Nm_SUM/N_SUM       hide
0 MIG_SAT = Nm_SAT/N_SAT       hide
0 MIG_MAL = Nm_MAL/N_MAL       hide
0 MIG_AMU = Nm_AMU/N_AMU       hide
```

*Fastsimcoal input files used to estimate the South Asian tiger population demography conditional on estimated demography of Asian tigers*

Note that fixed parameter values taken from the previous estimation are rescaled parameters that take into account the mutation rate assumed in these estimations (1e-8 per site per generation, instead of 0.35 e-8 per site per generation used to compute estimated values reported in Supplementary Tables 7 and 8).

TIGERS8e.tpl

```
//Simple model of tiger differentiation, no bottleneck
9  samples to simulate
//Population effective sizes
3766
N_META_IND
8960
6814
N_GHOST_ASIA
N_SAT_CI
N_SAT_NE
N_SAT_SA
N_SAT_SI
//Samples sizes
0
0
6
0
0
4
4
4
6
//Growth rates : negative growth implies population expansion
0
0
0
0
0
0
0
0
0
//Number of migration matrices : 0 implies no migration between demes
2
//Migration matrix 0
0 0 0 0 0 0 0 0 0
0 0 0 0 0 0 0 0 0
0 0 0 0 0 0 0 0 0
0 0 0 0 0 0 0 0 0
0 0 0 0 0 0 0 0 0
0 0 0 0 0 0 0 0 0
0 0 0 0 0 0 0 0 0
0 0 0 0 0 0 0 0 0
0 0 0 0 0 0 0 0 0
//Migration matrix 1
0 0 0 0 3.018842e-05 0 0 0 0
0 0 0 0 m_Asia2IND 0 0 0 0
0 0 0 0 1.321606e-06 0 0 0 0
0 0 0 0 2.993084e-05 0 0 0 0
0 0 0 0 0 0 0 0 0
0 m_SAT_CI   0 0 0 0 0 0 0
0 m_SAT_NE   0 0 0 0 0 0 0
0 m_SAT_SA   0 0 0 0 0 0 0
0 m_SAT_SI   0 0 0 0 0 0 0
//historical event: time, source, sink, migrants, new deme size, new growth rate, migration matrix index
26
T_RB 0 0 0 0.2034946 0 0 instbot  //Recent bottleneck in SUM
T_RB 2 2 0 0.1204263 0 0 instbot  //Recent bottleneck in MAL
T_RB 3 3 0 0.237151  0 0 instbot  //Recent bottleneck in AMU
T_RB 5 5 0 RB_SAT_CI  0 0 instbot             //Recent bottlenecks in INDIA
T_RB 6 6 0 RB_SAT_NE  0 0 instbot
T_RB 7 7 0 RB_SAT_SA  0 0 instbot
T_RB 8 8 0 0.3105766  0 0 instbot
T_RB 0 0 0 1 0 1
TD_SAT_CI  5 5 0 IB_SAT_CI   0 1 instbot //Old bottlenecks at divergence time in INDIA
TD_SAT_CI  5 1 1 1           0 1
TD_SAT_NE  6 6 0 IB_SAT_NE   0 1 instbot
TD_SAT_NE  6 1 1 1           0 1
TD_SAT_SA  7 7 0 IB_SAT_SA   0 1 instbot
TD_SAT_SA  7 1 1 1           0 1
TD_SAT_SI  8 8 0 IB_SAT_SI   0 1 instbot
TD_SAT_SI  8 1 1 1           0 1
TD_META_IND 1 1 0 IB_META_IND 0 1 instbot
TD_META_IND 1 4 1 1           0 1
640 0 0 0 0.124119     0 1 instbot   //Old bottleneck in SUM
640 0 4 1 1            0 1
544 2 2 0 0.003208539  0 1 instbot  //Old bottlenecks in MAL
544 2 4 1 1            0 1
529 3 3 0 0.01528991   0 1 instbot  //Old bottleneck in AMU
529 3 4 1 1            0 1
TDMAX 4 4 0 RES_ANC    0 0              //Ancestral population size change after last divergence
16397  4 4 0 1.068355  0 0   instbot    //Instantaneous bottleneck in ancestral population
//Number of independent loci [chromosome]
1 0
//Per chromosome: Number of contiguous linkage Block: a block is a set of contiguous loci
1
//per Block:data type, number of loci, per generation recombination and mutation rates and optional parameters
FREQ 1 0 1.0e-8 OUTEXP
```

TIGERS9B3Mv1_r8e.est

```
// Priors and rules file
// *********************

[PARAMETERS]
//#isInt? #name   #dist.#min  #max
//all Ns are in number of haploid individuals

//Population sizes -----------------------------------------
1 N_ANCEST         unif  1000  50000  output REFERENCE //Ancestral size (meta pop size as well)
1 N_META_IND       unif  5000  100000 output
1 N_SAT_CI         unif  1000   10000  output bounded
1 N_SAT_NE         unif  1000   10000  output bounded
1 N_SAT_SA         unif  1000   10000  output bounded
1 N_SAT_SI         unif  3924   3924  output bounded //Fixed to previously estimated value
1 N_GHOST_ASIA     unif  5797   5797  hide   //Fixed to previously estimated value
1 T_RB             unif  57     57   hide    //Fixed to rescaled time

//Divergence times -----------------------------------------
1 TD_META_IND      unif  100 900  output bounded
0 TD_REL_SAT_CI    unif  0   1       hide bounded
0 TD_REL_SAT_NE    unif  0   1       hide bounded
0 TD_REL_SAT_SA    unif  0   1       hide bounded
0 TD_REL_SAT_SI    unif  0   1       hide bounded

//Bottleneck intensities -----------------------------------
0 RB_SAT_CI        logunif  0.001 1  output
0 RB_SAT_NE        logunif  0.001 1  output
0 RB_SAT_SA        logunif  0.001 1  output
//0 RB_SAT_SI        logunif  0.001 1  output
0 IB_SAT_CI        logunif  0.001 1  output
0 IB_SAT_NE        logunif  0.001 1  output
0 IB_SAT_SA        logunif  0.001 1  output
0 IB_SAT_SI        logunif  0.001 1  output
0 IB_META_IND      logunif  0.001 1  output

//Gene flow rates ------------------------------------------
0 NM__Asia2IND     logunif 1e-4 5 output
0 NM__SAT_CI       logunif 1e-4 5 output
0 NM__SAT_NE       logunif 1e-4 5 output
0 NM__SAT_SA       logunif 1e-4 5 output
0 NM__SAT_SI       logunif 1e-4 5 output

[RULES]

[COMPLEX PARAMETERS]

//Divergence times ----------------------------------------
1 TDS_META_IND    =  TD_META_IND-T_RB hide
1 TD_PLUS_SAT_CI  =  TD_REL_SAT_CI  * TDS_META_IND hide
1 TD_PLUS_SAT_NE  =  TD_REL_SAT_NE  * TDS_META_IND hide
1 TD_PLUS_SAT_SA  =  TD_REL_SAT_SA  * TDS_META_IND hide
1 TD_PLUS_SAT_SI  =  TD_REL_SAT_SI  * TDS_META_IND hide

1 TD_SAT_CI  =  TD_PLUS_SAT_CI  + T_RB output
1 TD_SAT_NE  =  TD_PLUS_SAT_NE  + T_RB output
1 TD_SAT_SA  =  TD_PLUS_SAT_SA  + T_RB output
1 TD_SAT_SI  =  TD_PLUS_SAT_SI  + T_RB output
1 TDMAX      =  TD_META_IND %max% 829        output

//Migration rates ------------------------------------------
0 m_Asia2IND = NM__Asia2IND/N_META_IND hide
0 m_SAT_CI   = NM__SAT_CI/N_SAT_CI     hide
0 m_SAT_NE   = NM__SAT_NE/N_SAT_NE     hide
0 m_SAT_SA   = NM__SAT_SA/N_SAT_SA     hide
0 m_SAT_SI   = NM__SAT_SI/N_SAT_SI     hide

//Ancestral resize
0 RES_ANC    = N_ANCEST/N_GHOST_ASIA  hide
```
